# Loop Plasticity Drives Paralog-Specific Recognition in BET ET Domains

**DOI:** 10.1101/2025.11.11.687848

**Authors:** Guadalupe Alvarez, Elizabeth Sebastian, Arup Mondal, Monica J. Roth, Gaetano T. Montelione, Alberto Perez

## Abstract

The bromodomain and extraterminal domain (BET) family of proteins recognizes diverse peptide motifs via their conserved ET domains, yet exhibits paralog-specific binding preferences with unclear structural origins. Here, we use extensive molecular dynamics simulations and ensemble-based analyses to investigate how two known peptide epitope binders – one from the host regulatory protein NSD3 and one from the murine leukemia virus integrase – interact with the ET domains of BRD3 and BRD4.

Our results reveal that differences in peptide recognition are not driven by large conformational changes, but by subtle variations in loop dynamics and local secondary structure. Two divergent residues (positions 35 and 36) in the flexible loop connecting helices *α*2 and *α*3 shape the conformational ensemble of each paralog by modulating the formation of flanking helices (*η*1 and *η*2), which in turn control the opening of the peptide-binding cavity. These intrinsic dynamical differences influence the number and stability of accessible binding modes, with BRD3 showing greater plasticity and weaker binding, particularly for NSD3. Our findings support a model in which local sequence variation tunes conformational selection and induced fit mechanisms, offering a structural rationale for paralog-specific targeting of BET proteins.

**TOC Graphic:** 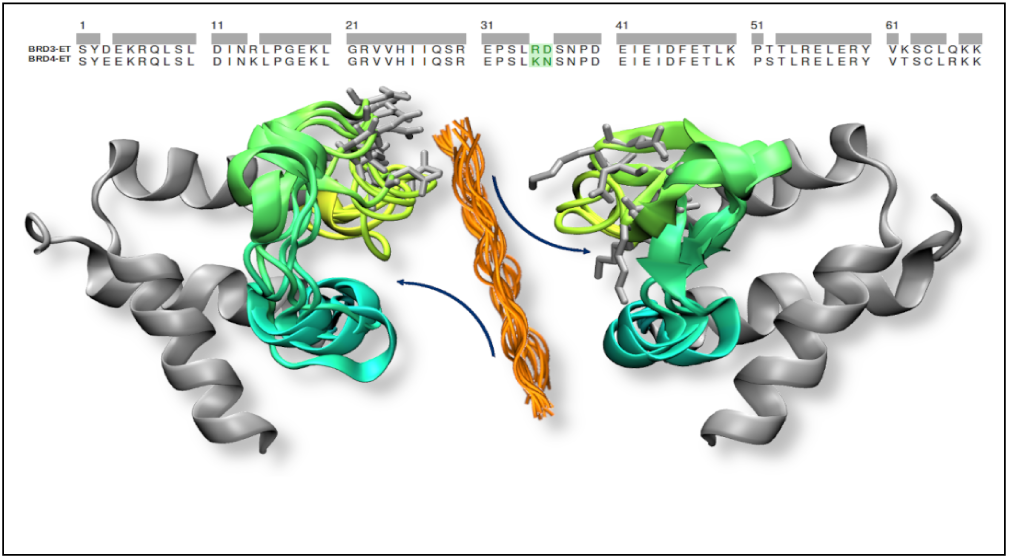

## Introduction

Bromodomain and Extra-Terminal (BET) proteins are key regulators of gene expression, particularly genes involved in immune responses^1,2^. Their misregulation has been linked to a range of diseases, including various cancers^3–6^. Humans express four BET paralogs—BRD2, BRD3, BRD4, and BRDT—which are differentially distributed across tissues and implicated in distinct pathological contexts^7^. Each BET protein can interact with multiple host and viral proteins, making them attractive targets for therapeutic intervention^7,8^.

All BET proteins contain at least two bromodomains (BD1 and BD2), which recognize acetylated lysine residues on chromatin, and a C-terminal extraterminal (ET) domain, which mediates interactions with regulatory and viral proteins. While bromodomains have been the primary focus of drug discovery efforts^9^, their high sequence homology has led to pan-inhibition, reduced specificity, drug resistance, and toxicity^10–13^. In contrast, the ET domain engages a diverse set of transcription factors, chromatin remodelers, and viral proteins through short linear peptide motifs ^14–19^. In this way, the ET domain functions as an interaction module, directly recruiting chromatin regulators and transcription factors, thereby integrating various signaling processes. This capacity to recruit effectors through diverse peptide motifs, which exhibit varying binding modes and affinities, suggests new opportunities for more targeted therapeutic strategies.

The ET domain itself is a compact, 68-residue three-helix bundle. It typically binds peptide epitopes located within intrinsically disordered regions of its partners, which fold into defined secondary structures upon binding. NMR structures of ET-peptide complexes reveal a striking diversity of binding modes, including helices, extended strands, and *β*-hairpins^18–21^. This plasticity is enabled in part by a flexible 20-residue loop between helices *α*2 and *α*3, which adapts its conformation depending on the bound peptide. Reported binding affinities span a broad range ^22^—from ∼ 100 *nM* to ∼ 950 *µM* —reflecting both the structural versatility of the ET domain and the sequence-dependent nature of its interactions. Although the ET domain is highly conserved across BET family members, some peptides exhibit preferential binding to specific paralogs. For instance, the previously characterized peptide epitopes of LANA and JMJD6 have been reported to bind BRD4-ET but not BRD3-ET^19,23^. However, our recent work has revealed that the ET domain can recognize previously uncharacterized binding motifs, including longer and structurally distinct epitopes that deviate from the canonical alternating charged/hydrophobic sequence pattern typically used to define ET-binding peptides^24^. Moreover, we have proposed alternative binding regions within the full-length LANA or JMJD6 proteins. Notably, one such motif has been experimentally validated, suggesting that prior reports may have overlooked biologically relevant interfaces or underestimated paralog-specific recognition. These findings underscore the need to reassess the molecular determinants of ET-mediated recognition and specificity and emphasize the importance of tools that can systematically explore the binding potential of flexible peptide motifs.

Recent advances in integrative modeling—such as MELD^22,25,26^ and AF-CBA^27^—have enabled the systematic discovery of novel ET-binding peptides by combining structural predictions with relative affinity ranking. These methods have already identified and helped validate new epitopes that deviate from canonical motifs, revealing binding modes previously unrecognized in experimental studies^24^. However, while these tools are powerful for epitope discovery, they are not yet sufficiently quantitative to resolve subtle energetic or structural differences that underlie paralog-specific selectivity. This limitation stems in part from the complexity of modeling folding-upon-binding events.

In contrast, conventional molecular dynamics (MD) simulations, when initiated from well-defined bound and unbound states, offer a complementary route to understand selectivity. By comparing the behavior of a particular peptide in complex with different ET paralogs, we can probe whether differences in binding modes, dynamics, or conformational plasticity of the receptor contribute to observed differences in affinity—without being confounded by the energetic cost of folding, which is presumed similar across complexes. Similarly, we can assess how different peptides interact with the same receptor, revealing sequence-dependent effects that may modulate binding strength or specificity.

In this study, we explore the molecular basis of paralog selectivity in peptide binding to BET ET domains. Using molecular dynamics (MD) simulations initiated from NMR-derived complex structures, we characterize the bound and unbound conformational ensembles of BRD3-ET and BRD4-ET. Specifically, we ask whether subtle sequence differences between paralogs translate into distinct structural or dynamic signatures that could influence peptide binding. By capturing changes in conformational plasticity across states and paralogs, we aim to shed light on the mechanistic underpinnings of ET-mediated selectivity.

## Methods

### Systems of study

We retrieved the seven existing NMR structures involving BRD3/4-ET from the PDB. BRD3-ET was found in its unbound form (PDB ID: 7JMY^21^) and binding four different peptides: BRG1 (PDB ID: 6BGH^20^), CHD4 (PDB ID: 6BGG^20^), TP (PDB ID: 7JQ8^21^), and NSD3 (PDB ID: 7JYN^21^). While there was no unbound structure for human BRD4-ET, there are two structures of complexes binding different peptides: JMJD6 (PDB ID: 6BNH^19^) and LANA (PDB ID: 2ND0^18^).

Out of the four BET proteins, structural efforts involving the ET domain have focused on BRD3 and BRD4 as they are the ones more relevant in disease. However, the experimental data does not describe these systems uniformly: only the BRD3-ET domain has been studied in its free form, and the complexes include different peptides for BRD3 and BRD4. The high degree of sequence and structural homology allowed us to model starting structures of the missing systems by mutating the seven residues that differ between BRD3-ET and BRD4-ET. We generate complementary sequences by rebuilding their structures based on backbone and *C_β_* coordinates using *tleap* ^28^. This approach allowed us to model BRD3-ET in complex with LANA and JMJD6, and BRD4-ET in complex with CHD4, BRG1, NSD3, and TP. We also modeled the unbound BRD4-ET state by removing LANA and JMJD6 from their respective NMR complex structures. After energy minimization of these systems, the resulting equilibrated MD ensembles quickly lost memory of the starting conformations and adopted features consistent with the unbound state.

### Molecular Dynamics protocol

All the simulations were performed using the AMBER 22 biomolecular simulation suit^29^. Each system was set up using the Amber ff19SB protein forcefield^30^ and the OPC water forcefield^31^ with explicit solvent in an octahedral solvent box of 10 Å. Na and Cl ions were added to neutralize the charges. The energy of the systems was minimized over six steps. The initial steps applied restrictions to the protein that were gradually removed. Each step involved 4000 cycles of minimization implementing steepest descent followed by conjugate gradient. The systems were then heated to 298.15 K using a Langevin thermostat with a collision frequency of 1 *ps^−^*^1^ for 1 ns, followed by a NPT equilibration using a Berendsen barostat for 10 ns. After equilibration, each simulation was run in a NPT ensemble at 298.15 K using a Langevin thermostat and a Monte Carlo barostat. Hydrogen mass repartition^32^ and SHAKE^33^ algorithm were implemented to reach a time step of 4 fs. We ran simulations in triplicate for 7.2 *µ*s each, saving coordinates every 100 ps. Thus for each system we had an accumulated 21.6 *µ*s sampling time.

### Analysis

MD trajectory analysis was performed using CPPTRAJ^34^, along with custom Python and R scripts. Meta-ensembles were constructed by combining three replicates per system; in some cases, trajectories from different peptide complexes were merged to define consensus receptor ensembles for PCA. Conformational space was analyzed via principal component analysis (PCA) using backbone atoms of residues 6–61. An average structure from the concatenated trajectories defined the reference frame, and eigenvectors were computed from the covariance matrix. Kernel density estimation (KDE) was used to project PC1 and PC2 distributions, from which free energy surfaces were estimated as Δ*G* = −*k_B_T* log *P_i_*. Representative conformations were extracted via k-means clustering (k = 10), by implementation of MDAnalysis^35^ and Scikit-learn^36^ combined in a Python script, with RMSD as clustering metric. Free energy profiles along PC1 and PC2 were compared using Kullback–Leibler divergence^37^.

To complement PCA, we used ELViM (Energy Landscape Visualization Method)^38^, a multidimensional projection based on pairwise structural dissimilarities, applied to the same backbone region. Trajectories from both BRD3-ET and BRD4-ET, in bound and unbound states, were combined into a single meta-ensemble using a common reference topology. Additional analysis details are provided in the Supplementary Methods.

## Results

### Loop rearrangements differentiate BRD3 and BRD4 binding behavior

The BRD3 and BRD4-ET domains differ by seven residues (see panel A in Figure 1), yet adopt the same 3-helix bundle fold in NMR structures of their peptide-bound complexes. While there is an NMR structure of the unbound BRD3-ET domain in solution (PDB ID 7JMY^21^), there is no structure for the unbound BRD4-ET domain. Notably, none of the seven mutations are directly involved in forming the hydrophobic binding pocket (see panel B in Figure 1). Interestingly, many of these changes are conservative in nature (e.g. D3E, R14K, R35K, T52S).

**Figure 1:**
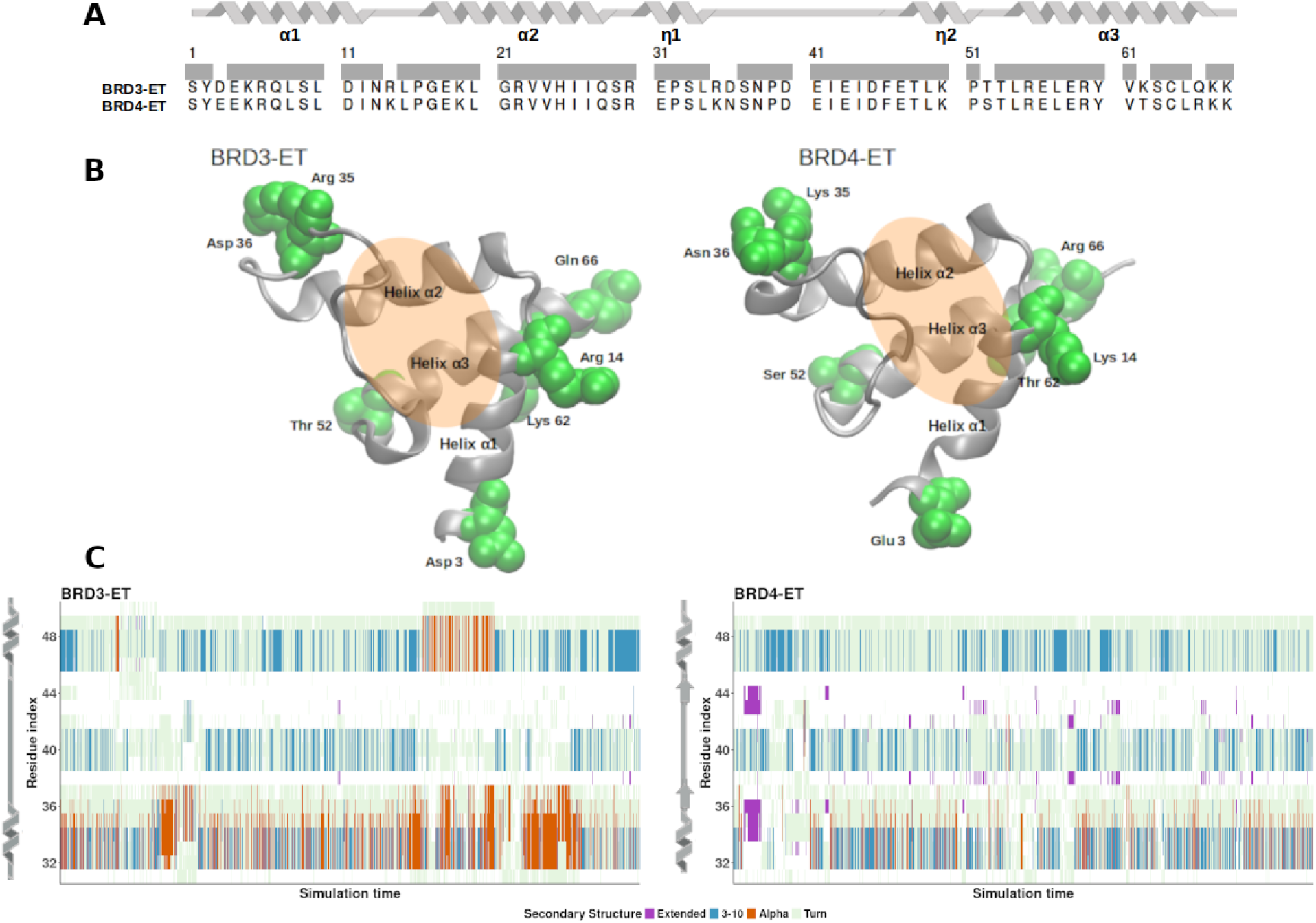
A. Sequences of the globular region of BRD3-ET (residues 573 to 640 in the BRD3 sequence) and BRD4-ET (residues 611 to 678 in the BRD4 sequence) used for the MD simulations. The gray bar highlights the sequence identity. The panel also shows the secondary structure elements of the ET domains, generated with the software SSDraw^39^. B. Representation of the initial structures used for MD simulations of BRD3-ET (left) and BRD4-ET (right) in their unbound form. Mutated residues are highlighted in green and the orange ellipse schematically represents the peptide-binding pocket. C. Secondary structure content calculated for the residues of the *α*2 − *α*3 loop region as function of the simulation time from the MD, for the unbound forms of BRD3 and BRD4-ET. The figure also highlights the differences observed in secondary structure features of each paralog along the simulations.

The main difference between the bound and unbound forms are residues in the loop region connecting helices *α*2 and *α*3 (expanding residues 31 to 50). Upon peptide binding, this loop undergoes significant conformational changes driven by a zipper-like interaction interface alternating hydrophobic and charged residues. Four central residues in this loop form key interactions with peptide side chains, including two conserved negatively charged residues (Glu43 and Asp45) that engage positively charged complementary residues on the peptides.

Flanking these core residues, both the N- and C-terminal regions of the loop exhibit secondary structure preferences. In particular, the N-terminal part adopts a short helical structure (*η*1 helix) in some NMR structures and our simulations show that this helix has a transient nature. Two of the seven BRD3/4-ET sequence differences are located in this *η*1 helix region. On the C-terminal side, our simulations reveal a less frequent helix or turn-like feature (*η*2). Additionally, BRD4-ET occasionally samples a short *β*-hairpin conformation in this region – a feature not observed in BRD3-ET (Figure 1C).

In the sections that follow, we examine how the intrinsic structural preferences of the BRD3- and BRD4-ET loops—particularly in the *η*1, central contact, and *η*2 regions—influence their dynamics and peptide-binding behavior.

### Peptides that bind as hairpins occupy the full ET pocket and form stable complexes

We analyzed six known ET-binding peptides based on published NMR structures: four with BRD3 (BRG1, CHD4, TP, NSD3) and two with BRD4 (LANA, JMJD6). The peptides are derived from both viral (LANA, TP) and host ( NSD3, BRG1, CHD4, JMJD6) proteins and display a range of conformations — extended single strands (CHD4, BRG1, LANA), *β*-hairpins (NSD3, TP), and an *α*-helix (JMJD6) (the sequences and Uniprot ID of the peptides are provided in SI Table 1). To assess their behavior and possible determinants of selectivity, we performed triplicate molecular dynamics simulations for each peptide–ET complex with both BRD3 and BRD4. While the ET domain remained stable in terms of backbone RMSD, some peptides dissociated during the simulations (see Fig. SI 1).

**Table 1:** KL divergence (PC1 and PC2) between ensemble distributions. Cells are color-coded by the larger of the two values: blue < 0.5 (similar), gray 0.5–1.5 (moderate), red *>* 1.5 (divergent).

| Comparison Type | Systems Compared | KLD PC1 | KLD PC2 | Interpretation |
| --- | --- | --- | --- | --- |
| Binding-induced<br>(within BRD3) | unbound vs TP | 0.7 / 1.31 | 0.34 / 0.48 | Moderate conformational change with binding |
|  | unbound vs NSD3 | 0.17 / 0.76 | 0.26 / 0.27 |  |
| Binding-induced<br>(within BRD4) | unbound vs TP | 2.00 / 1.55 | 1.64 / 1.47 | Large conformational change with binding |
|  | unbound vs NSD3 | 2.72 / 0.93 | 1.61 / 1.09 |  |
| Paralog difference<br>(same state) | unbound BRD3 vs BRD4 | 0.2 / 0.85 | 0.17 / 0.09 | Unbound BRD3 and BRD4 are similar<br>Moderate to large differences upon binding |
|  | TP bound BRD3 vs BRD4 | 1.16 / 0.29 | 0.28 / 0.29 |  |
|  | NSD3 bound BRD3 vs BRD4 | 2.3 / 0.58 | 0.3 / 0.31 |  |
| Peptide difference<br>(same paralog) | BRD3-TP vs BRD3-NSD3 | 0.5 / 0.38 | 0.08 / 0.08 | TP and NSD3 induce similar ensembles within each paralog |
|  | BRD4-TP vs BRD4-NSD3 | 0.17 / 0.15 | 0.07 / 0.03 |  |

LANA and JMJD6—both among the weakest reported binders—dissociated rapidly. BRG1 and CHD4, which bind with moderate affinity (*K_d_*= 7 ± 1 *µ*M and 95 ± 10 *µ*M, respectively^20^), remained bound longer but still showed fluctuations and unbinding events in some replicates. In contrast, TP and NSD3 peptides remained bound throughout all simulations with both BRD3 and BRD4. TP, the highest affinity binder in the set (*K_d_* = 13±10 nM at 10*^◦^*C, 90 ± 10 nM at 25*^◦^*C)^22^ forms a stable hairpin-like interface. While the NSD3 epitope used in this study is a weaker binder (*K_d_* = 250 ± 150 *µ*M at 10*^◦^*C)^22^ compared to BRG1 and CHD4, it spans the entire ET-binding pocket—unlike the other two—and adopts a stable hairpin conformation in our simulations. These observations highlight that stable ET binding in MD correlates with peptides adopting a *β*-hairpin conformation that fully spans the binding pocket.

MD simulations are known to sometimes over-stabilize compact structures, particularly for intrinsically disordered proteins^40–42^. In contrast, the observed unbinding of LANA and JMJD6—despite this known MD bias—underscores the labile nature of these complexes. The persistent binding of NSD3 and TP instead suggests the formation of energetically favorable, well-matched interfaces.

Given their persistent binding in simulations, we focus the remainder of this study on the TP and NSD3 peptides. As previously noted, both peptides bind the BRD3-ET domain as *β*-hairpins, but they do so in opposite orientations: TP binds in a C-to-N-terminarl direc-tion (spanning *β*7*^′^* to *β*6*^′^*), while NSD3 binds N-to-C-terminal (from *β*1*^′^* to *β*2*^′^*), following the nomenclature of Aiyer et al. ^21^. These orientations align alternating hydrophobic and positively charged residues on the peptides with complementary hydrophobic and negatively charged residues within the ET loop. Notably, the loop regions of the peptides occupy different positions relative to the ET loop—TP’s hairpin loop aligns near the N-terminal end of the ET loop, while NSD3’s aligns near the C-terminal end.

Figure SI 2 illustrates these two complexes, showing the secondary structure element nomenclature and peptide sequences. Despite adopting similar hairpin structures, the opposite orientations suggest that the ET domain accommodates diverse binding modes. This raises the possibility that paralog-specific selectivity may arise not only from sequence differences, but also from differences in the dynamic response of ET to distinct binding modes. To investigate this, we used experimentally determined NMR structures of BRD3-ET complexes and homology-modeled BRD4-ET complexes to compare conformational dynamics and binding mechanisms across paralogs.

### Comparative conformational dynamics of BRD3 and BRD4

The NMR structures already point at a significant rearrangement of the loop region connecting helices *α*2 and *α*3. To characterize conformational differences between BRD3 and BRD4 in unbound and peptide-bound states, we applied Principal Component Analysis (PCA) and the Energy Landscape Visualization Method (ELViM). While ELViM provides a reference-free mapping of the ensembles, PCA projections require eigenvectors derived from a chosen reference set. In systems with subtle conformational changes, such as those present in these systems, the choice of reference can significantly affect interpretability.

We evaluated four PCA reference ensembles: unbound systems (*unbound REF* ), systems bound to the Tail Peptide (*TP REF* ), systems bound to NSD3 (*NSD*3 *REF* ), and a combined set of all TP- and NSD3-bound systems across BRD3 and BRD4 (*TP NSD*3 *REF* ). As shown in Figure SI 3, the reference choice impacts the shapes of the projected ensemble distributions. While *unbound REF* highlights global contraction upon binding, the resulting distributions are diffuse and less informative for distinguishing between bound states. In con-trast, the *TP NSD*3 *REF* yields a more discriminative projection, revealing key differences across paralogs and peptide complexes (Figure 2).

**Figure 2:**
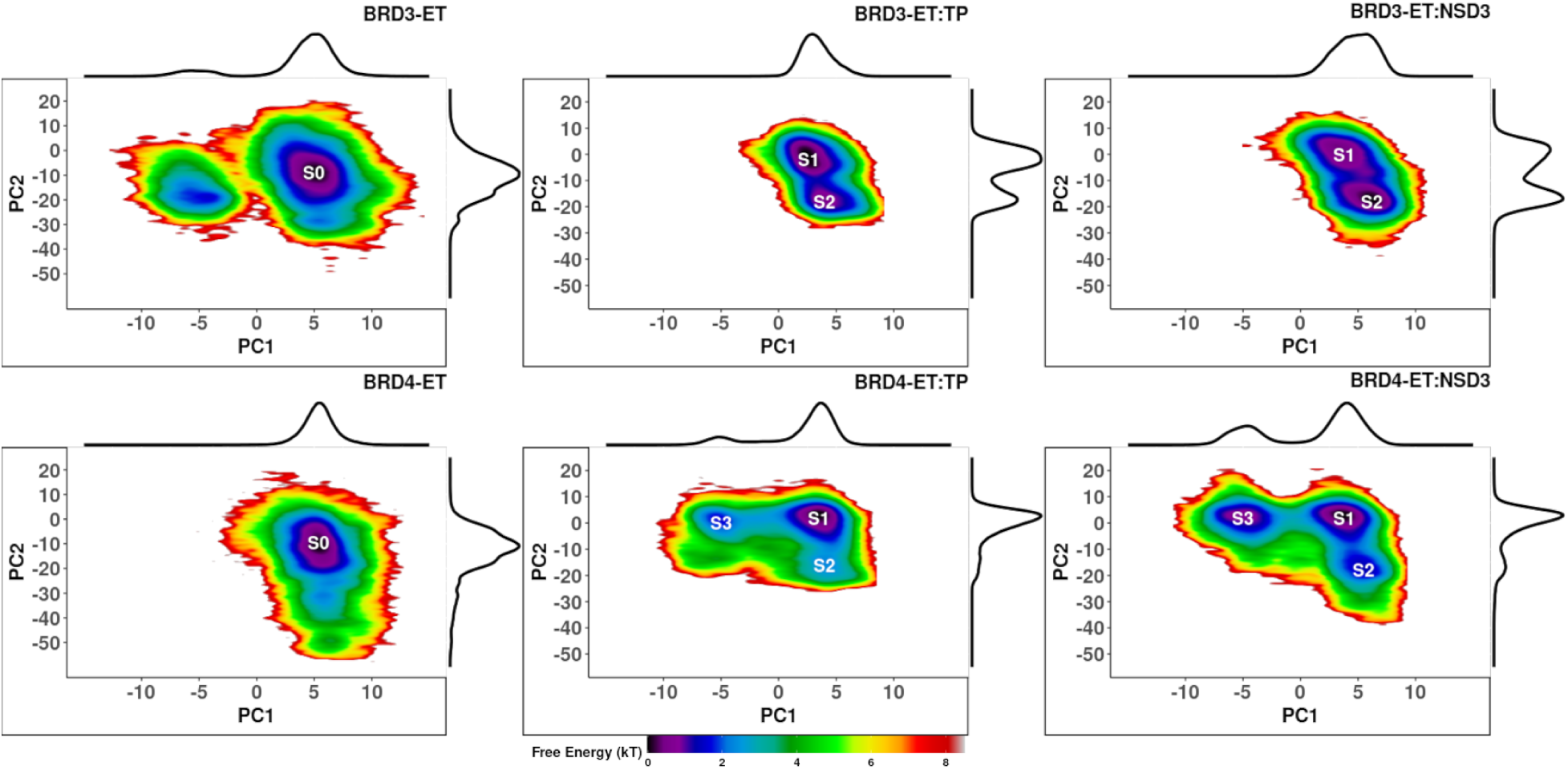
PCA using the *TP NSD*3 *REF* reference frame. The PC and the projections were calculated considering the backbones of the residues 6 to 61. Estimation of the free energy (*k_B_T* ) in the first two principal components (PC1 and PC2) space, calculated with KDE, for BRD3-ET (top row) and BRD4-ET (bottom row) unbound and bound forms. Each panel also shows the marginal distribution of PC1 and PC2. The metastable states are highlighted as *S*0, *S*1, *S*2 and *S*3.

We therefore adopted *TP NSD*3 *REF* as our standard reference for PCA. This choice enabled us to identify marked differences between BRD3 and BRD4, both in their unbound states and in their conformational response to peptide binding (see Fig. 2). Both BRD3 and BRD4 sample a similar core in the unbound state (*S*0), but each also accesses minor distinct regions. Upon binding, both paralogs undergo conformational shifts to populate new metastable states. Two metastable states (*S*1 and *S*2) are common to both BRD3-ET and BRD4-ET in their bound forms, while a third state (*S*3) is unique to BRD4-ET.

Although bound BRD4 samples three metastable states, its ensemble is dominated by *S*1 across both peptides. In contrast, BRD3 samples two states with comparable populations, with the dominant state (defined by integrating the population in that state) depending on the bound peptide (*S*1 for TP, *S*2 for NSD3). PC1 captures a transition between *S*1 and *S*3, a pathway sampled by BRD4-ET but not accessed by bound BRD3-ET. PC2 is related to describing a transition between metastable states *S*1 and *S*2.

Free energy estimates projected along PC1 and PC2 (Fig. SI 4) show that while the metastable minima occupy broadly similar regions across systems, the BRD4-bound ensembles exhibit higher energy barriers between states, suggesting more restricted interconversion. In contrast, BRD3-ET displays a unique but broader basin along PC1, and a low free energy barrier along PC2 separates states *S*1 and *S*2, facilitating frequent transitions between them. For NSD3 in particular, shallow barriers and well-defined alternate minima are observed in the BRD3 ensemble. Overall, these energy landscape features suggest that deeper energetic basins may be associated with reduced conformational flexibility, while broader landscapes with accessible sub-states may underlie greater plasticity in binding modes.

To further quantify ensemble differences, we computed the Kullback–Leibler (KL) divergence along the first two principal components (Table 1; see also Figs. SI 5 to SI 7). As expected, bound *versus* unbound comparisons yield high KL divergence, reflecting peptide-induced conformational shifts. By contrast, divergence between TP- and NSD3-bound forms is low within each paralog, indicating that both peptides induce broadly similar conformational preferences. Divergence across paralogs reveals that BRD3 and BRD4 are more similar to each other in their unbound form than in any of their peptide-bound states. Notably, asymmetries in KLD along PC1 suggest that BRD4-ET samples a metastable state in the bound form that is never accessed by BRD3-ET.

While these results highlight structural responses to binding, they do not unambiguously explain binding specificity or affinity and their origin. Both paralogs exhibit a core region that is similar in their unbound state. However, each paralog samples additional substrates that are orthogonal to each other: BRD4-ET extends sampling along regions in the second principal component, whereas BRD3-ET samples additionally along the first principal component. However, upon binding, BRD4-ET samples more metastable states with *S*1 as the core metastable state, independently of which peptide binds. Conversely, BRD3-ET only samples two metastable states, but with closer populations indicating both are significantly sampled in the ensemble, but which one is the preferred metastable state depends on which peptide binds (*S*1 for TP and *S*2 for NSD3). Further details about each ensemble’s principal components are depicted as porcupine plots in Figures SI 8-10.

We additionally explore the differences between the metastable states by extraction of representative structures from dominant PCA clusters. For the unbound forms (Fig. SI 11 and SI 12), we observe differences in secondary structure propensities—such as a higher prevalence of *β*-hairpins in BRD4 and helix-turn motifs in BRD3—and in the spatial relationship between the ET loop and the adjacent helices that define the peptide-binding cavity, particularly with respect helix *α*1. We also observe loss of structure in the C-terminal loop. Upon peptide binding (Fig. SI 13 and SI 14), major structural rearrangements compared to the unbound forms, are observed in the loop and in the adjacent short helix: *η*2 region in the BRD3-ET complexes and *η*1 region in the BRD4-ET complexes. However, differences are also observed in the *α*1-helix when superimposing structures to the unbound forms. This structural differences observed in bounded forms are consistent with the residual dynamics observed in NMR and chemical shift perturbation studies,^21^ comparing bounded and unbounded states of BRD3-ET.

### Complementary visualization of the conformational space with ELViM

Because PCA relies on a chosen reference ensemble and may miss subtle structural heterogeneity, we complemented it with ELViM^38^. ELViM projects conformational ensembles into a low-dimensional space based solely on pairwise structural dissimilarities, offering a reference-free and topology-aware view of structural variability. While PCA highlights major transitions between discrete metastable states, ELViM offers a finer-grained view of the structural landscape, revealing additional substructure within those basins and exposing subtle differences in conformational entropy between paralogs and bound forms. We applied ELViM to a metaensemble comprising both unbound and peptide-bound forms of BRD3 and BRD4. Figure 3 shows the resulting projections, with free energies estimated via kernel density estimation (KDE) and mapped onto a consistent color scale across all systems.

**Figure 3:**
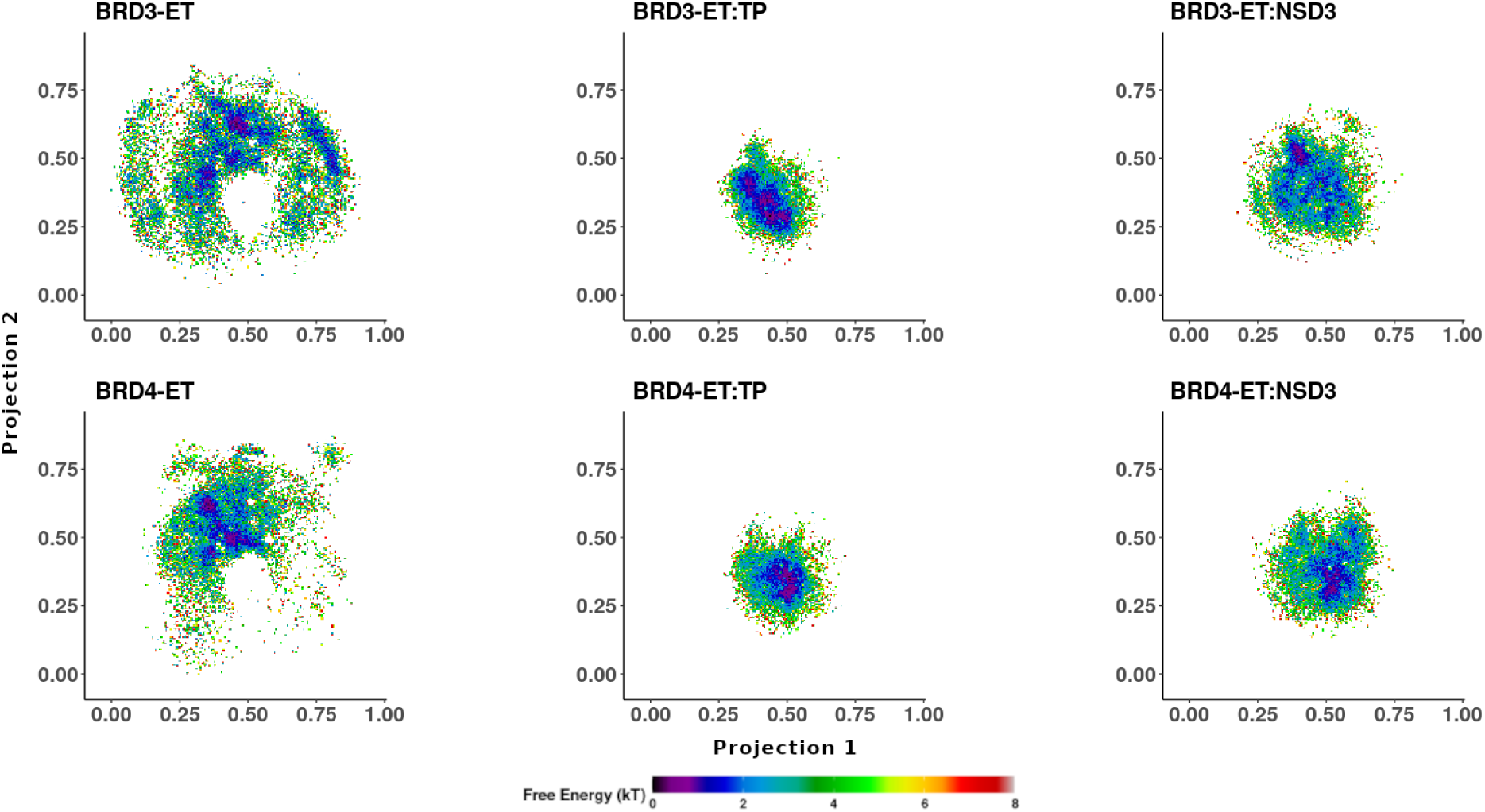
ELViM projections of BRD3 and BRD4-ET ensembles, with free energy estimated via KDE. All maps are scaled identically. Axes are included for reference but do not correspond to physical coordinates; structural relationships are encoded in pairwise distances. Each panel reflects the conformational landscape of one system.

The unbound ensembles of BRD3 and BRD4 show broad and overlapping distributions, with BRD3 exploring a wider conformational space. BRD4-ET samples a compact region with three closely related metastable states, which also appear in BRD3-ET but at lower occupancy. In addition, BRD3-ET accesses two distinct high-population regions not sampled by BRD4.

Peptide binding markedly compacts the conformational space for both paralogs, shifting populations into defined low-energy regions with minimal overlap with their unbound ensembles. This compaction reflects both a restriction in conformational space and a peptide-induced structural rearrangement—likely involving stabilization of the ET loop region that forms part of the binding cavity. TP binding induces greater compaction than NSD3, in both paralogs.

The effect of peptide binding differs between BRD3 and BRD4. For BRD4, both TP and NSD3 lead to tight ensembles with two main metastable states located in similar regions. In contrast, BRD3 samples different regions depending on the bound peptide: TP binding yields three distinct low-energy basins, whereas NSD3 binding results in a more diffuse distribution with many low-occupancy states and a single dominant one. Interestingly, most of the states sampled by BRD3:TP, BRD4:TP, and BRD4:NSD3 are also accessible to BRD3:NSD3, but with low probability. The dominant state in BRD3:NSD3 is unique to that system, suggesting a distinct conformational preference not observed elsewhere. This broader view suggests that *S*1 and *S*2 are composed of multiple sub-basins, revealing subtle conformational heterogeneity that is not resolved in PCA alone.

Representative structures extracted from ELViM maps for the unbound forms (Fig. SI 15 and SI 16) also point to differences in secondary structure propensities of the loop region between paralogs, in the relative positioning of the *α*1-helix, and in the folding of the C-terminal loop. Mainly, these structural differences suggest different tendencies across paralogs toward pre-organization of the loop prior to peptide binding, which could contribute to the observed differences in peptide binding preferences or affinities.

### The ***α***2–***α***3 Loop as a Determinant of Paralog-Specific Peptide Recognition

Global conformational analyses across BRD3 and BRD4 reveal that differences in peptide binding and selectivity trace back to structural and dynamical features of the *α*2-*α*3 loop. This loop displays the greatest variability in flexibility, secondary structure propensity, and dynamic correlations, and its stabilization upon peptide binding differs depending on both the peptide and the receptor paralog.

To quantify these differences, we analyzed structural and dynamic features of the loop across unbound and bound forms. First, we examined the loop’s overall structural variability via RMSD distributions. We first computed RMSD distributions for the full ET domain and for the loop region (residues 31–50) in the presence and absence of peptides (Fig. 4A). The unbound BRD3 ensemble shows broader, often bimodal RMSD distributions, reflecting greater conformational diversity. In contrast, BRD4 displays narrower distributions with a long tail. Peptide binding stabilizes the loop in both paralogs, with TP inducing more pronounced narrowing of distributions than NSD3. This stabilization is especially evident in BRD4, which maintains a narrow, unimodal distribution upon binding, consistent with a preorganized binding interface.

**Figure 4:**
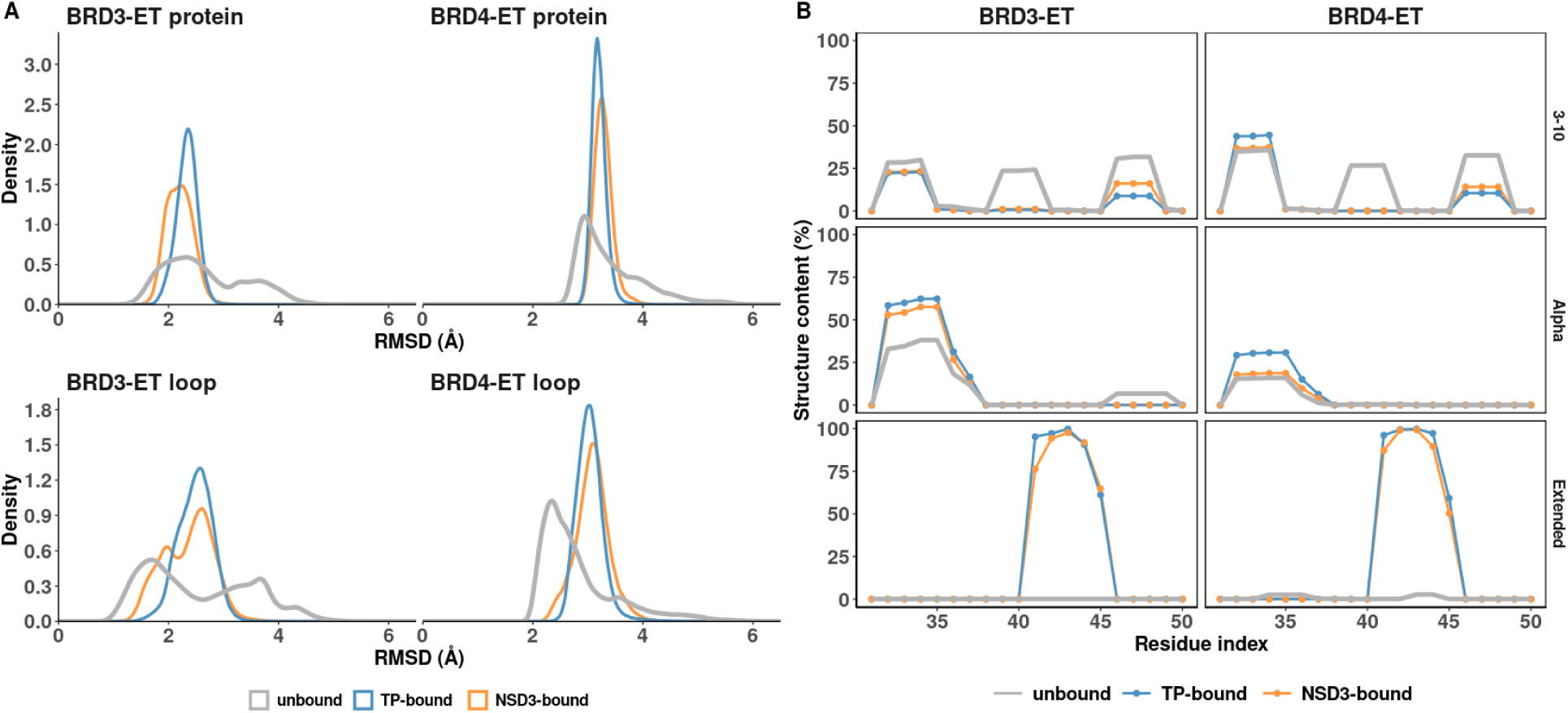
A. RMSD distributions. On the top, we considered the protein receptor, residues 6 to 61 against the reference structure. On the bottom, we only considered the loop region, between residues 31 to 50 against the reference structure. B. Secondary structure content of the α2 − α3 loop region, represented as distributions, of key structural features.

These RMSD findings align with secondary structure analysis (see the distribution in Fig. 4B and the time series in Fig. SI 17). In the presence of peptides, residues 40–46 in the loop consistently form an extended *β*-strand, matching the interaction motif of alternating charged and hydrophobic residues. Flanking this strand, short helices form transiently: residues 31–41 (*η*1) and 45–51 (*η*2). BRD3 exhibits more frequent helix formation in the unbound form, especially in *η*1, and this helicity increases upon binding, while BRD4 has a shorter *η*1 helix, found mostly as a 3-10 helix, with small changes upon binding. The *η*2 region interconverts between a 3-10 helix and a loop, with helix population decreasing upon binding in both paralogs. Interestingly, as mentioned previously and shown in Fig. 1C, BRD4 presents a very small hairpin population in its unbound form, where one of the strands overlaps with the region that becomes a strand upon peptide binding. These subtle differences in helix and strand propensities along the loop are related to the effect and rearrangement upon binding, as well as the tightness of the binding. The formation and disruption of these helices during our simulations indicate a possible mechanism for opening/closing the binding site.

To complement RMSD-based stability assessments, we next analyzed local fluctuations via Root mean square fluctuation (RMSF) analysis (Fig. SI 18). RMSF confirms the *α*2–*α*3 loop as the most flexible region in the unbound forms (residues 26–51). Peptide binding reduces fluctuation in the central loop region (residues 40–46), while flexibility remains higher in the flanking segments (residues 30–40), particularly around positions 35 and 36—two sites that differ between BRD3 (R35, D36) and BRD4 (K35, N36). TP binding suppresses loop flexibility more effectively than NSD3. Furthermore, a sensitivity analysis (Fig. SI 19) highlights the importance of residues 35 and 36. Interestingly, unbound BRD3 does not showcase this behavior, which only arises upon binding. On the contrary, BRD4-ET shows this behavior in its unbound form and is further accentuated during binding.

Normal mode analysis provides additional insight into the motion patterns. BRD4-ET exhibits more collective motions of the binding site, with coordinated shifts of residues 30–41 that modulate access to the cavity. In contrast, BRD3-ET displays a hinge-like tilting motion involving the loop and *α*1-helix (residues 11 and 35), suggesting alternative conformational states that could accommodate peptides differently. These motions are captured by eigen-vectors 2 and 3, which describe the opening/closing of the ET cavity. Notably, this motion is accompanied by transient formation of surface pockets enriched in polar and charged residues. These pockets may represent cryptic allosteric sites capable of modulating the open/closed equilibrium of the binding site, and thus merit future exploration (see Fig. SI 20).

Correlated motion matrices further support distinct intrinsic dynamics (Fig. 5). BRD3 and BRD4 display markedly different correlation fingerprints upon peptide binding. Notably, the correlation pattern is conserved across peptides for a given receptor—suggesting that the loop’s dynamic response is encoded in the receptor, not the peptide. This points to an intrinsic paralog-specific mechanism for shaping binding outcomes.

**Figure 5:**
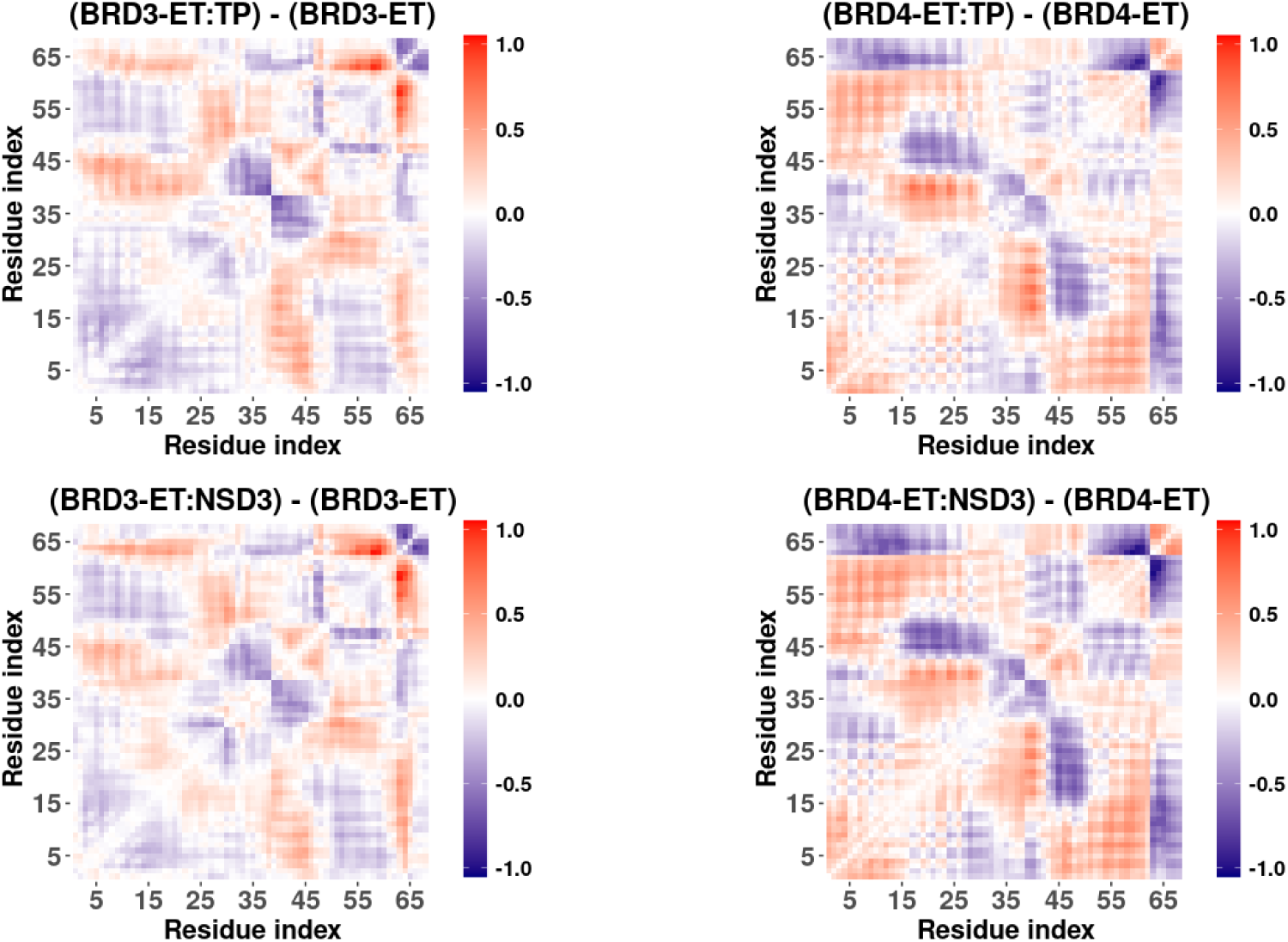
Difference correlation matrices for BRD3-ET and BRD4-ET upon peptide binding. Each panel shows the pairwise correlation difference between the peptide bound-system and the unbound system for BRD3 and BRD4-ET and for each peptide.

In line with this, analysis of peptide location reveal that BRD4 enforces a more narrow distribution of the position of the peptide in the binding site TP binds both paralogs in a well-defined mode, while NSD3 shows broader positional distributions—especially in BRD3, which supports two distinct binding modes (Fig. 6).A Kolmogorov-Smirnov statistical test comparing the RMSD distribution for each peptide bound to BRD3 and BRD4 yielded p-values smaller than 0.05, confirming that observed differences in the location of the peptide for each receptor are statistically significant. This suggest that BRD4 promotes a more constrained binding mode, while BRD3 exhibit greater plasticity, allowing weaker binders as NSD3 to explore multiple binding modes. We propose that BRD3 may favor a conformational selection mechanism for these weaker binders, whereas the more constrained BRD4 binding may be related to the loop reorganization and consistent with induced fit mechanism.

**Figure 6:**
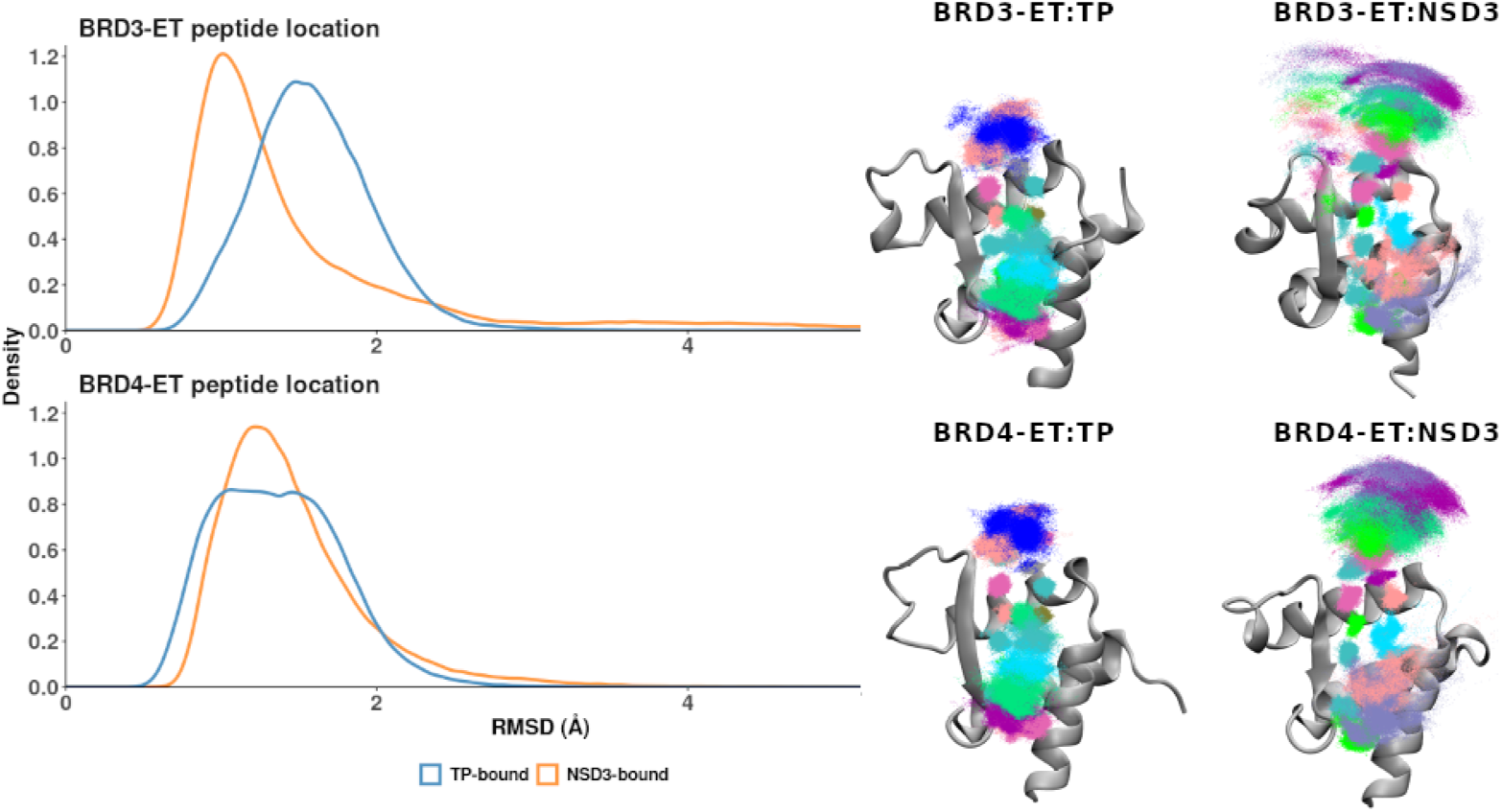
RMSD distribution of peptide localization and representation of the peptide ensembles. The RMSD represents peptide location against the aligned protein receptor, it was calculated excluding the disordered regions of each peptide, considering an average structure as reference but without fitting. In the structure illustrations, each dot represents a superimposed frame of the peptide, aligned to the protein receptor and the different colors represent different residues in the peptide.

The complexes may be further stabilized by interactions beyond the loop. NSD3 has been reported to interact with the hydrophobic pocket of BRD3-ET through a phenylalanine residue (*Phe18* )^21,43^. Our simulations are in agreement with these observations, since this *Phe* residue makes close and frequent contacts (*<* 5 Å) with several residues on ET domain *α*2. Interestingly, our TP-bounded simulations reveal an analog interaction between a tryptophan residue (Trp5) and residues on ET *α*2—helix (Fig. SI 21). We explore these interactions further using a per-residue interaction decomposition with the MM/PBSA approach^44^. This analysis acknowledges that beyond the typical positive charged and hydrophobic interactions, the most favorable contributions came from this *Phe* for NSD3 and the *Trp* residue for TP, with the contribution being stronger for TP (Fig. SI 21). These contacts highlight a secondary anchor point in the binding mechanism and may underlie TP’s higher binding affinity.

### The Role of ***η***1/***η***2 in Binding-Site Regulation

Coupled with its flexibility, the loop propensity to form transient helical turns and small helices (*η*1 and *η*2) is fundamental for peptide binding. To characterize these regions beyond secondary structure, we looked at *i* → *i* + 4 distance distributions along the loop region in both paralogs as well as in the presence/absence of peptides (see Figs. SI 22 and SI 23). Bimodal distributions in some of these regions reflect changes between hydrogen-bonded N-H…O atoms involved and helices and conformations in extended states.

The *η*2 region (residues 44–50) exhibits the most consistent behavior regarding helix formation. In their unbound form, distributions are broad and multimodal. Binding to either peptide in either paralog produces narrow, unimodal distributions without significantly shifting the location of the peaks, indicating a stabilization of pre-existing local structures. The mid-loop region (residues 40–46) also goes from broad distributions to narrow distributions upon binding – but now the peaks of the distributions shift towards extended conformations compatible with the formation of a *β*-strand required for binding the peptides. As this extended conformation is not sampled in the unbound form, there is little overlap between bound and unbound distributions, and a similar behavior is observed for both paralogs.

Most striking is the *η*1 region (residues 31–38), which includes the two BRD3/BRD4 mutations at positions 35 and 36. In BRD3, these residues (R35, D36) form a salt bridge, both in the unbound form and in the complexes (see Figure SI 24), an interaction that is lost in BRD4 (K35, N36). This region shows high conformational heterogeneity in the unbound forms and retains bimodal or broad distributions even after binding—especially in BRD3:NSD3 (Figure 7.A). These distributions reflect helical-to-extended conformational switching. In BRD3, peptide binding populates conformations absent in the unbound state. The *η*1 N-terminal region exhibits narrower distributions, likely due to the presence of the salt bridge, while the C-terminal region and beginning of the loop, show less-defined distributions. This increased conformational flexibility and broader loop opening in BRD3 may explain the presence of multiple peptide binding modes and lower binding affinity. The two paralogs show similar behavior in the unbound form – but differential behavior in the bound form, where the location of maxima is in the same places, but their populations vary considerably, in particular for distances 32-36, 33-37, and 36-40 (see Figure 7.A).

**Figure 7:**
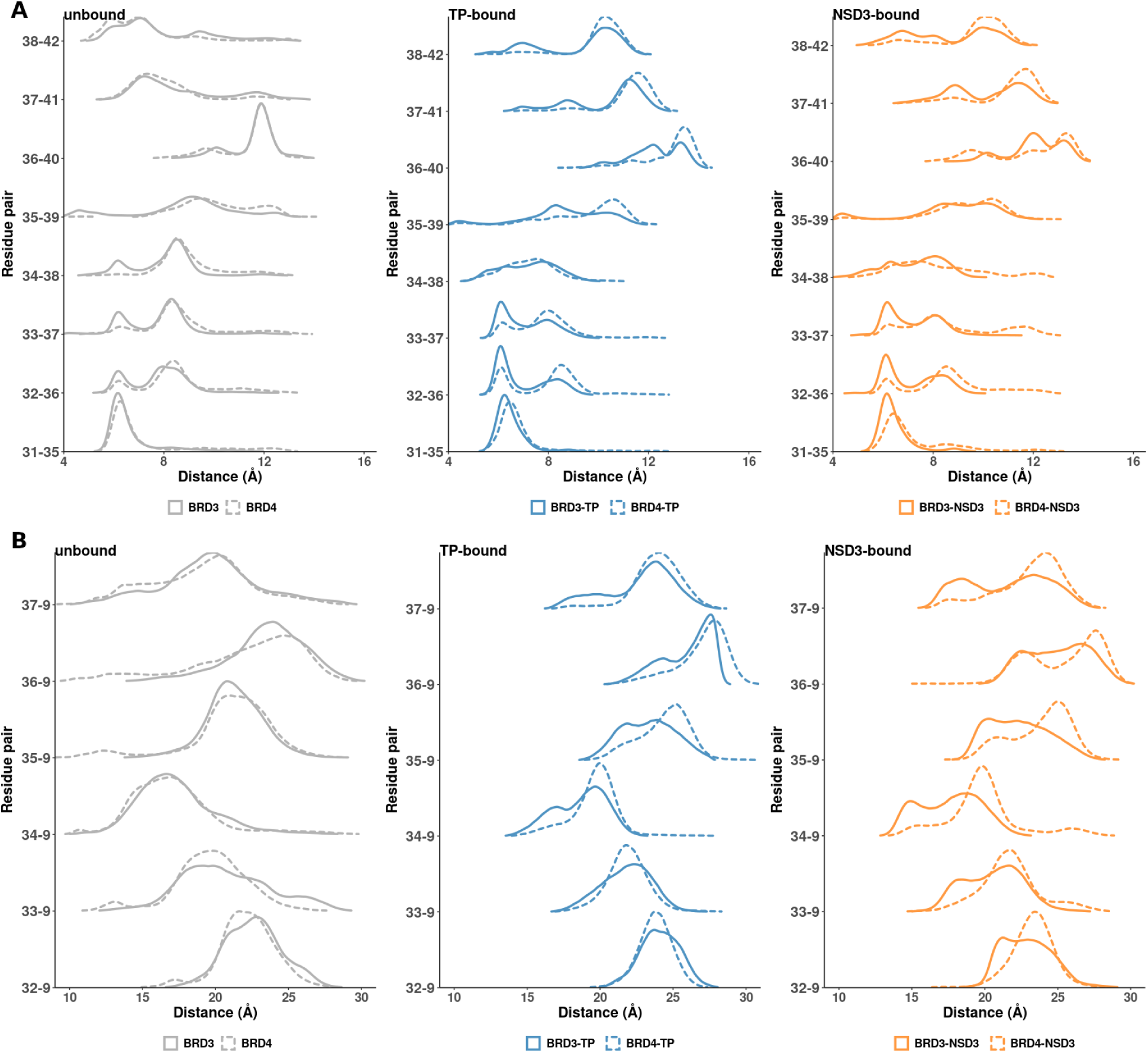
A. Distances i −→ i+4 between the *C_alpha_* of loop residues that define the η1-helix and exhibit larger differences between the unbound and bound forms of BRD3 and BRD4-ET. B. Distances between the *C_beta_* of loop residues that define the η1-helix and the *C_beta_* of residue 9 located in α1-helix.

To further dissect how these changes affect binding site opening regulation, we analyzed distances between loop regions (*η*1, *β*, and *η*2) and helix *α*1, which together define the peptide-binding cavity (Figs. SI 25–SI 27).

Near the *η*2 region (Fig. SI 27) we see *η*2 − *α*1 distances becoming unimodal narrow distributions consistent with stabilizing the size of the binding site. This is also observed for the *β* − *α*1 region (see Fig. SI 26). In both cases there are no appreciable differences in the distributions for both paralogs, and the narrowing of distributions is consistent with maintaining an open binding site when a peptide is bound.

The *η*1 − *α*1 region again show the largest differences upon binding (see Fig SI 25 and some particular selected distances in Figure 7.B). While the unbound distributions are nearly identical across paralogs, peptide binding induces marked changes in both the location of the maxima and the shape of the distributions. The nature of the peptide makes especially pronounced differences for BRD3-ET. This peptide-dependent heterogeneity is most prominent around residues 35 and 36 in the NSD3-bound BRD3 ensemble, underscoring their role in modulating loop opening. Variability in binding site opening may permit alternative peptide binding modes. Since BRD3 has a natural tendency towards heterogeneity, the specific nature of peptide interactions, especially those residues binding between the *η*1 helix and *α*1 region, appears to influence both binding affinity and binding mode. For example, TP’s loop region includes a tryptophan residue that forms stable interactions with BRD3, stabilizing a dominant binding mode. Similarly, NSD3 contains a Phenylalanine residue that is positioned to perform the same role as the Tryptophan in TP, but might lack an equal level of stabilization, given the difference observed in the peptide ensembles.

Together, these results reinforce the role of *η*1 and *η*2 helices as modulators of loop conformation and binding-site accessibility. Mutations in *η*1 (residues 35/36) expand the conformational ensemble of BRD3, enabling greater plasticity in peptide recognition, particularly for weak binders like NSD3. This dynamic behavior is tightly coupled to the formation of strand structure in the core binding region and supports a model in which loop opening/closing motions—governed by flanking helix formation—gate access to the ET binding cleft.

Overall, these results support a model in which the *α*2–*α*3 loop acts as a dynamic and structural switch that tunes binding specificity and stability across BET paralogs. Although BRD3 and BRD4 exhibit similar behavior in the unbound state, their bound conformations diverge markedly. Despite their high sequence similarity, the two paralogs differ in conformational plasticity and induced motions, leading to distinct interaction landscapes with peptide partners. In particular, peptide binding produces a larger shift in conformational distributions for BRD3-ET than for BRD4-ET.

## Discussion

BET proteins are critical regulators of gene expression and chromatin remodeling. They are also hijacked by viral proteins such as Murine Leukemia Virus Integrase (MLV) and Kaposi’s Sarcoma Herpes Virus Latency Associated Nuclear Antigen (LANA), which exploit the ET domain to interact with host chromatin machinery^45,46^. However, the mechanisms underpinning these interactions—particularly how selectivity arises among the highly conserved BET paralogs—remain poorly understood.

In this study, we used molecular dynamics simulations to probe the structural and dynamical landscapes of BRD3 and BRD4 ET domains in their unbound forms and when bound to two distinct peptides: the MLV integrase tail peptide (TP) and a segment of the host regulatory protein NSD3. These peptides share a common sequence motif of alternating charged and hydrophobic residues, which forms *β*-strand-like interactions with the ET domain loop.

Our simulations demonstrate that complex stability is strongly influenced by peptide length and the ability to fully occupy the binding site. Although alternative NSD3 epitopes have been proposed^24^, raising questions about which is biologically relevant, the construct used in this study spans the entire ET-binding pocket, was characterized by NMR, and remained stable in our simulations. Moreover, it exhibits a higher experimental binding affinity than shorter variants (12–13 residues), consistent with the requirement of occupying the whole binding site^18,20,21^.

Our results highlight that differences in binding behavior between peptides and BET paralogs arise not from large structural rearrangements, but from subtle differences in loop dynamics, secondary structure propensities, and correlated motion. Specifically, the *α*2–*α*3 loop emerges as the primary modulator of binding-site accessibility and stability. Despite high overall sequence conservation, two divergent residues (positions 35 and 36) in this loop region lead to substantial differences in structural flexibility, helix/strand formation, and peptide binding modes.

Across multiple metrics—RMSD, RMSF, secondary structure content, PCA/ELViM projections, and essential dynamics—the BRD4-ET domain exhibits tighter, more stable ensembles when bound to peptides, particularly TP. BRD3, by contrast, shows broader distributions and a greater number of accessible metastable states, particularly when binding the weaker NSD3 peptide. Conformational changes upon binding are not restricted to loop region, they propagate across BRD3 and BRD4 domains, consistent with our observation of chemical shift perturbations throughout the BDR3 ET domain upon TP or NSD3 complex formation^22^. This suggests a higher degree of conformational plasticity in BRD3 and supports a model of conformational selection for weak binders, versus induced fit for strong binders like TP.

Importantly, our results show that loop dynamics are not just passive outcomes of binding, but active regulators of binding site shape. We propose that short helices flanking the binding region (*η*1 and *η*2) act as tunable hinges that regulate opening and closing of the peptide-binding groove. Subtle sequence differences affect the frequency and stability of these secondary structure elements, thereby shifting the ensemble toward open or closed states. These dynamics are sufficiently distinct between BRD3 and BRD4 to suggest paralog-specific structural preferences—even in the absence of large conformational changes.

This dynamic mechanism also explains the existence of multiple binding modes for NSD3 in BRD3, contrasted with a single, more compact binding mode in BRD4. Our peptide ensemble analysis further confirms that BRD4 restricts the conformational space of bound peptides more tightly than BRD3, especially for TP. A tryptophan residue in TP interacts with the *α*2-helix in both receptors, but contact strength and geometry may be modulated by loop preorganization. Interestingly, some of these contact residues are among the mutated positions between paralogs, which may contribute to differences in binding affinity.

Although binding affinity values reported in the literature vary widely depending on peptide length and experimental conditions, trends in our simulation data align with observed affinities: tighter peptide ensembles and reduced flexibility are associated with stronger binding (e.g., TP), while broader ensembles and persistent flexibility correlate with weaker binding (e.g., NSD3).

Altogether, our study suggests that:

1. Small sequence differences—particularly at positions 35 and 36—amplify into global dynamical changes that modulate binding behavior.
2. The *α*2–*α*3 loop is a key conformational switch that regulates access and stability of peptide interactions.
3. BRD3 and BRD4 possess distinct dynamical ”fingerprints” that persist regardless of which peptide is bound, opening the door for paralog-selective targeting.
4. The *η*1/*η*2 motif and the loop region more broadly could represent cryptic allosteric or druggable pockets, especially given their transient closure and cooperative motions. The differences in dynamic behavior open the door for paralog-specific targeting.

In sum, binding selectivity appears to arise not from differences in global structure, but from subtle allosteric regulation mediated by loop helices that act as hinges. These modulate access to the core binding groove through peptide-specific and paralog-specific reorganization of a conserved conformational landscape. This has implications for both functional understanding and therapeutic design. Future work will aim to identify peptides or small molecules that preferentially stabilize or disrupt specific conformations of BRD3 or BRD4, exploiting these dynamic differences for selective targeting.

We focused our study on BRD3 and BRD4-ET paralogs, but we hypothesized that the helices flanking the loop might also influence the loop dynamics of the other paralogs. BRD2-ET presents a close sequence similarity to BRD3-ET, while BRDT-ET is the most divergent. Particularly, residues 35 and 36 are conserved between BRD2 and BRD3-ET, but in BRDT the mutation in the position 35 (R in BRD3 *vs.* S in BRDT) is non-conservative regarding charge. These sequence differences are interesting to explore in future research, as they might influence the loop dynamic and binding specificity in the other paralogs.

## Conclusion

Our study reveals that the *α*2–*α*3 loop of the BET ET domain acts as a dynamic regulatory element that governs paralog-specific peptide recognition. Subtle sequence differ-ences—particularly at positions 35 and 36—profoundly influence loop flexibility, local secondary structure formation, and binding-site accessibility. These effects shift the conformational preferences of BRD3 and BRD4 and modulate how each paralog engages peptide ligands such as NSD3 and TP. Our findings support a model in which the *η*1 and *η*2 helices function as dynamic hinges that regulate access to a strand-binding interface via mechanisms of conformational selection or induced fit—distinctions that will require further investigation. Importantly, these dynamical signatures and the overall conformational landscape remain paralog-specific regardless of the bound peptide, providing a mechanistic basis for selective engagement. Beyond advancing our understanding of BET domain recognition, this work offers a structural and dynamic framework for designing paralog-selective therapeutics that exploit intrinsic differences within this conserved epigenetic reader family.

## Acknowledgement

The authors thank the support from the National Institutes of Health grant R35-GM141818 (to G.T.M.), R35GM122518 (to M.J.R.), and R01-GM149646 (to A.P.).

## Data availability

Molecular dynamic trajectories, corresponding to the dry concatenated triplicates, saved each 15 frames for storage purposes, and representative structures from PCA-based clustering presented in Figures SI 11 to SI 14, are available in the Zenodo repository under DOI 10.5281/zenodo.17418909.

