## Supplementary Methods and Figures for "Loop Plasticity Drives Paralog-Specific Recognition in BET ET Domains"

#### Evaluation of systems global stability

We evaluated the global stability of the 12 initial systems throughout the MD simulations calculating the backbone RMSD for the ET-domain (residues 6 to 61) (Fig. SI 1, top) using an average structure for each system obtained after concatenation of the triplicates.

We observed unbinding events during the simulations of the complexes with peptides that do not fully occupy the ET pocket. To characterize these events, we calculate the distances between the center of mass of the ET-loop (residues 30 to 50) and the center of mass of the peptide region containing the alternate positive charged and hydrophobic residues (Fig. SI 1, bottom). These peptide fragments were selected to include the main interaction regions and to exclude the flexible tails.

Table SI 1 summarizes the peptides used and their sequences, highlighting the regions known to interact with ET-domain.

SI Table 1: Peptides reported to form complexes to the ET-domain used in this work. The fragments of the sequences highlighted in bold correspond to the interaction regions.

| Peptide acronym | Sequence | Length | UniprotID (residue range) | Origin of the peptide |
|---|---|---|---|---|
| BRG1 | RSVK**VKIKL**GRK | 12 | P51532-1 (1591–1602) | Brahma-Related Gene 1 |
| CHD4 | KVAP**LKIKL**GGF | 12 | Q14839-1 (290–301) | Chromodomain Helicase DNA-binding protein 4 |
| JMJD6 | KWTLER**LKRK**YRN | 13 | Q6NYC1-1 (84–96) | Jumonji domain-containing protein 6 |
| LANA | NLQSSI**VKFK**KPLP TQPG | 19 | Q9QR71 (1098–1116) | Latency-Associated Nuclear Antigen of Kaposi's Sarcoma Herpesvirus |
| NSD3 | PE**IKLKI**TKTIQNG RELFESSLCGDLLN EVQASE | 34 | Q9BZ95-1 (151–184) | Histone-lysine N-methyltransferase Nuclear Receptor binding SET domain protein 3 |
| TP | SRLTWRVQRSQNP **LKIRL**TREAP | 23 | P03355 (1716–1738) | C-terminal tail peptide of Murine Leukemia Virus Integrase |

#### NMR structure of BRD3-ET:TP and BRD3-ET:NSD3 complexes and secondary structure nomenclature

The NSD3 an TP epitopes are part of a larger protein, where the peptide epitopes belong to disorder regions that fold upon binding. Accordingly, the nomenclature[1] for when they stabilize hairpins in their binding mode to ET corresponds to the order of the secondary structure elements they have in the complete sequence (see Fig. SI 2). We track secondary structure changes along the sequence using Cpptraj's DSSP[2] implementation to assign each residue of each trajectory frame a specific secondary structure. For simulations with a bound peptide we analyze the whole system even if we only report specific residues in the receptor afterwards.

#### PCA analysis for the different reference frames

As mentioned in the Main Text, the choice of reference has a significant impact on the shape and spread of the distributions. In Figure SI 3 we show the PCA obtained for bounded systems, considering the four reference frames. In general, the *unbound_REF* allows the visualization of the contraction of the conformational space upon binding. However, compared to the other references, the resulting distributions are broader and more diffuse. The bounded references yield more collapsed distributions, suggesting that the systems become more conformationally restricted. While $TP\_REF$ produces compact distributions, the $NSD3\_REF$ and $TP\_NSD3\_REF$ reveal signs of conformational separation or clustering, with some systems presenting multiple populated regions. Depending on the chosen reference frame, the conformational ensemble may appear compact or stretched, which can bias the interpretation of similarities and differences between systems. Therefore, we conclude that it is important to define more than one reference frame and explain how it was built.

##### Estimation of Free Energy difference between BRD3-ET and BRD4-ET along the first two PCs

To complement the ensemble comparison along a shared principal component (PC) frame shown in Figure 2 of the Main Text for the bound forms of BRD3-ET and BRD4-ET, we estimated the free energy profiles along each PC (see Fig. SI 4). We first computed the population distributions ($P_i$) along PC1 and PC2 using kernel density estimation (KDE), then derived the corresponding free energy profiles using $\Delta G = -k_B T \log(P_i)$. The resulting curves were referenced to the global minimum of the BRD3-ET TP-bound profile along PC1 (indicated by a dotted horizontal line) to enable comparison of relative energy barriers across systems.

##### Kullback-Leiber divergence

To quantify the similarities and differences of the distribution of PC1 and PC2, calculated for the different systems under the $TP\_NSD3\_REF$, we calculated the KL divergence[3] values for pairs of distributions. Figures SI 5, 6, 7 show the compared distributions along with the corresponding KL divergence values.

##### Normal Modes Representation

We visualize the first principal components (or normal modes) with the NMWizard[4] complement for VMD[5]. Figures SI 8, 9, 10 show porcupine plots capturing the first three principal components of each system. For the calculations, the frames were aligned to each system's reference structure, and the backbone for residues 6 to 61 was considered.

##### Representative structures from PCA and ELViM

We extracted representative structures performing a k-means clustering (with k = 10) in the space defined by the projection of the trajectories onto the first two PC of the $TP\_NSD3\_REF$. The clustering was performed by implementation of MDAnalysis[6] and

Scikit-learn[7] combined in a Python script, with RMSD as clustering metric. Figures SI 11 to SI 14 show representative clusters based on PCA projections. To better visualize conformational changes, we aligned the centroid structure of each cluster against the centroid of the top cluster of its corresponding unbound form and calculate the RMSD per residue. The resulting RMSD profiles were projected onto the structure representations to color them.

Figures SI 15 and SI 16 show representative structures extracted manually from the ELViM[8] projections. Since there were no cluster populations in this case, we selected one of the extracted structures to use as reference to calculate the RMSD per residue of the other structures against it. The resulting RMSD profiles were used to color the representations and visualize conformational changes.

When comparing the representative structure of the clusters obtained for each receptor, the main differences are observed in the loop region and in the N- and C-terminal regions. In the case of BRD3, we can observe some loss of secondary structure of the C-terminal region, involving residues 59 to 68 (centroid of cluster C5). In the case of BRD4, we observed that the loop can adopt a harping conformation (centroid of cluster C3) that appears to adopt an extended and well-defined conformation (centroid of cluster C5). The differences in the loop region are also observed in the representative structures extracted from the ELViM maps.

##### Fluctuation and perturbation analysis

We evaluate local flexibility by calculating RMSF profiles, considering the backbones of residues 6 to 61 (see Fig. SI 18). The profiles obtained for BRD3 and BRD4-ET are very similar. We observe the overall rigidization of the ET domain upon peptide binding. For the bounded forms of both ET domains, the loop region exhibits higher flexibility and, in particular, the maximum peaks are located around residues 35 to 37, which are part of the small $\eta$ helix.

We also perform a Perturbation-Response scan (PRS) adapted from reference,[9] to identify the residues that are the most sensitive to perturbations (see Fig. SI 19). In this analysis, a force is applied systematically to the alpha carbon of each residue. Then, the resulting displacement is calculated from the force and the covariance matrix obtained from the MD simulation. The resulting PRS matrix is then normalized by dividing each element of a column by the diagonal element in the same column. Row $i$ in the normalized matrix represents how a perturbation at residue $i$ affects each residue $j$, generating an "influence profile" of residue $i$ on the other residues. Column $j$, in turn, describes the response of residue $j$ to the perturbation of each residue $i$, representing a "sensitivity profile" of residue $j$.

The residues that are the most responsive after peptide binding are Ser37 for BRD3 and Asp36 for BRD4. Surprisingly, the loop region of the unbounded form of BRD4 is more sensitive than BRD3. We can relate this to the different propensity of the protein receptors to form helices.

##### Exploration of cryptic pockets in BRD3-ET simulations

Observation of the BRD3-ET simulations reveals a transient pocket between $\alpha 2$ and $\alpha 3$-helices (see Fig. SI 20). This observation suggests the potential existence of a cryptic pocket, which could be further explored as a novel druggable site. To reinforce our observations, we perform a pocket prediction with the software *fpocket*[10] for selected frames. The transient pocket that we observe between $\alpha 2$ and $\alpha 3$-helices has the highest *fpocket* score, which means that it is likely to be a site where small molecules could bind.

##### Binding interface analysis

To explore the interactions between BRD3-ET and BRD4-ET and the peptides, we combine a contact map with a per-residue decomposition of the binding free energy. We perform an MM/PBSA calculation[11] using the Ameber implementation[12] for both ET-domain receptors (see Fig. SI 21). Although it is acknowledged that MM/PBSA is not a precise method for absolute free energy calculations, especially because we did not consider the entropy contribution due to its high computational cost, it can be valuable for comparative analysis of similar binders. Here, we implemented the method to identify the interaction hot-spots in the peptide ligands, evaluating their relative contribution to binding free energy, recognizing that the flexibility of the unbound peptides is a limitation. For this calculation, we consider the electrostatic, van der Waals, and polar solvation terms to the free energy contribution and use it to complement the structural and dynamic information obtained in the contact map.

The contact map, as well as the MM/PBSA per-residue decomposition, show the distinct binding orientations of TP and NSD3 peptides upon binding. For TP, the most frequent contacts are residues 13 to 20 ($\beta 7'$ strand), and for NSD3, residues 3 to 9 ($\beta 1'$ strand), which are located in the binding interface with the ET loop. In both peptides, these regions contain the alternate positively charged and hydrophobic residues and exhibit the most favorable contribution to the MM/PBSA free energy. In contrast, the peptide residues that form the third strand in the $\beta$-hairpin upon binding are less frequent contacts and their energetic contribution is lower. Interestingly, for TP, the tryptophan (Trp5) residue presents the most favorable MM/PBSA contribution within the $\beta 6'$ strand. For NSD3, the phenylalanine (Phe18) residue located in the $\beta 2'$ strand present the most favorable contribution. This Phe residue of NSD3 has been reported to interact with the hydrophobic cleft of BRD3-ET[1,13]. These observations may support the importance of these residues in the stabilization of their respectively complexes with the ET-domains.

### Supplementary Figures

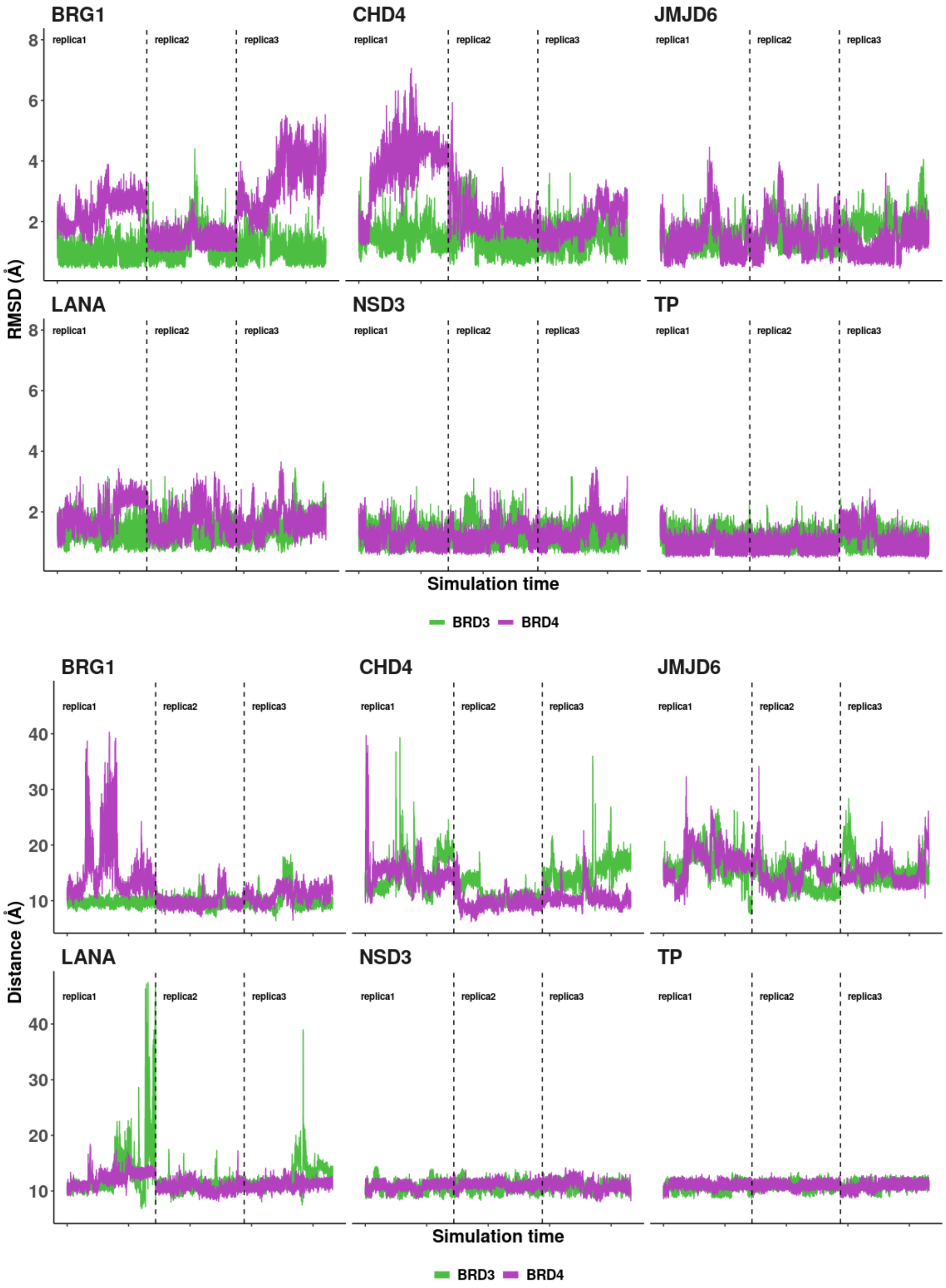


SI Figure 1: Top: Backbone RMSD of the ET-domain (residues 6 to 61) along the simulation time. The structures were aligned against and average structure of each system, after concatenation of triplicates. Bottom: Distance between the center of mass of the ET-domain loop (residues 30 to 50) and the center of mass of the peptide region that interacts with the ET-domain (see SI Table 1).

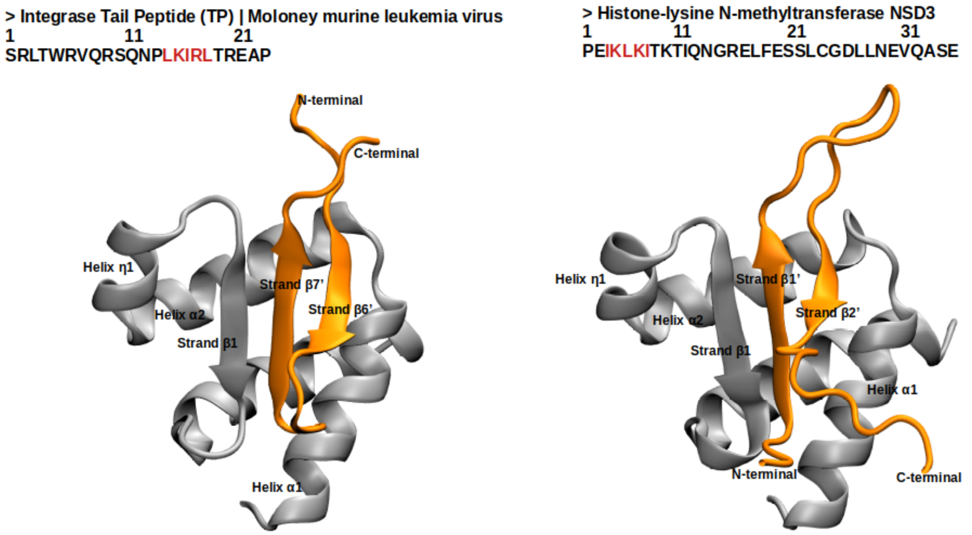


SI Figure 2: Left: sequence of the TP peptide and representation of the NMR derived structure of the complex with BRD3-ET (PDB ID 7JQ8). Right: sequence of the NSD3 peptide and representation of the NMR-derived structure of the complex with BRD3-ET (PDB ID 7JYN). The highlighted amino acids in the peptide sequences correspond to the alternate hydrophobic and positively charged residues.

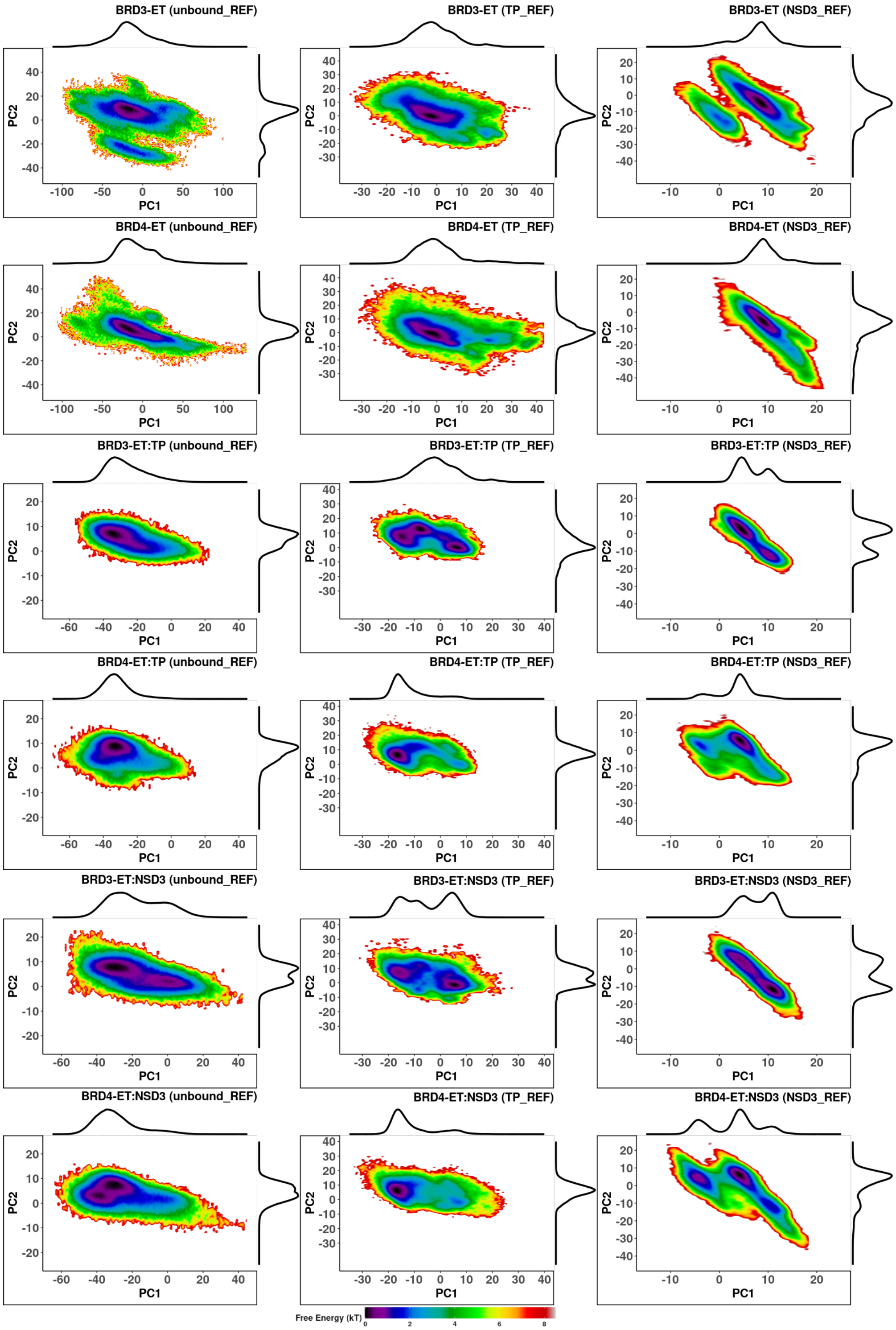


SI Figure 3: Estimation of the free energy ($k_BT$) and marginal distributions for the bounded systems onto the PC1 and PC2 of the different reference frames. The PC and the projections were calculated considering the backbones of residues 6 to 61. The first two rows show the BRD3 and BRD4-ET unbound form. The third and fourth rows show the BRD3 and BRD4-ET TP-bound forms, respectively. Finally, the last two rows show the BRD3 and BRD4-ET NSD3-bound forms, respectively.

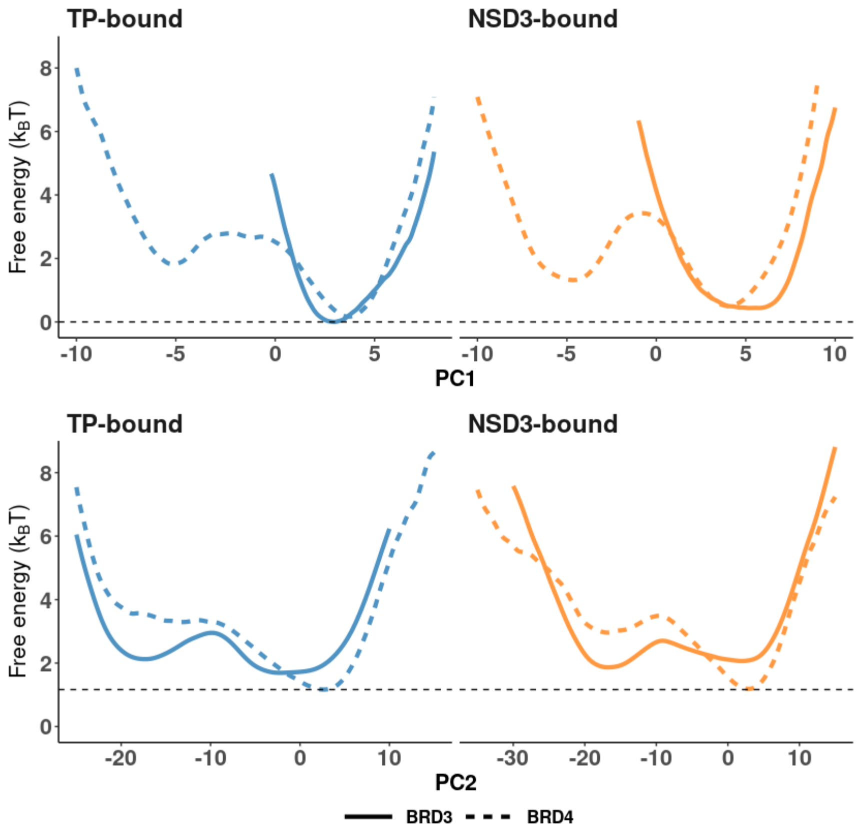


SI Figure 4: Estimation of Free Energy profile across PC1 (top) and PC2 (bottom).

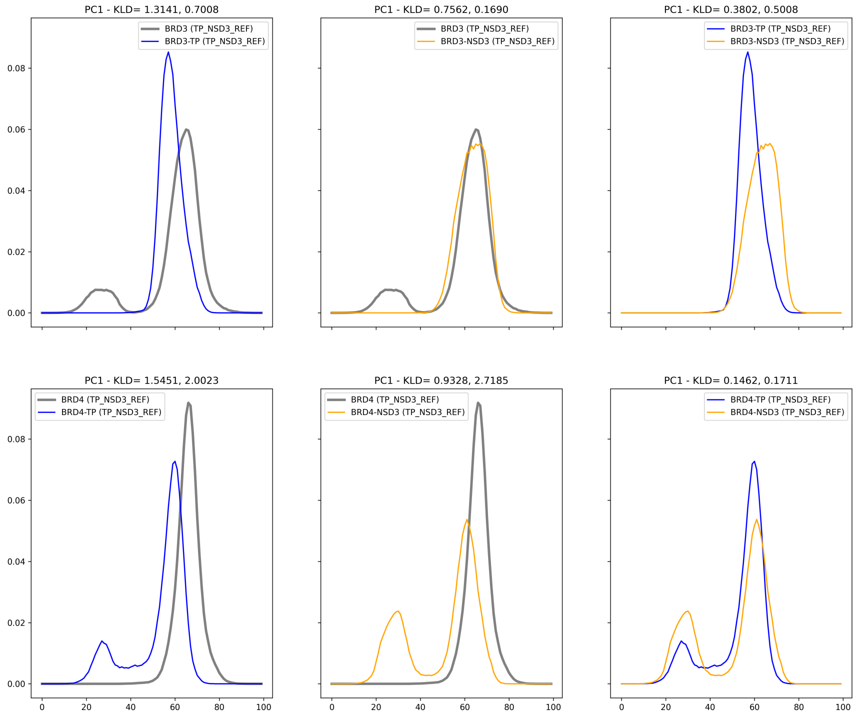


SI Figure 5: Distribution of the first PC and KL divergence values. Comparison between unbound and bound forms of each paralog (BRD3-ET on the top row and BRD4-ET on the bottom row).

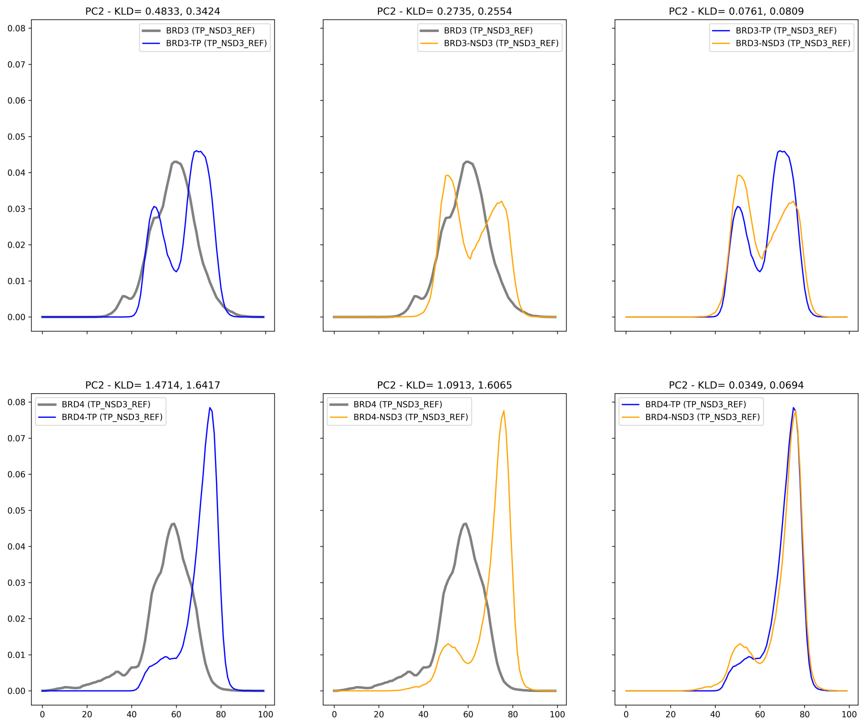


SI Figure 6: Distribution of the second PC and KL divergence values. Comparison between unbound and bound forms of each paralog (BRD3-ET on the top row and BRD4-ET on the bottom row).

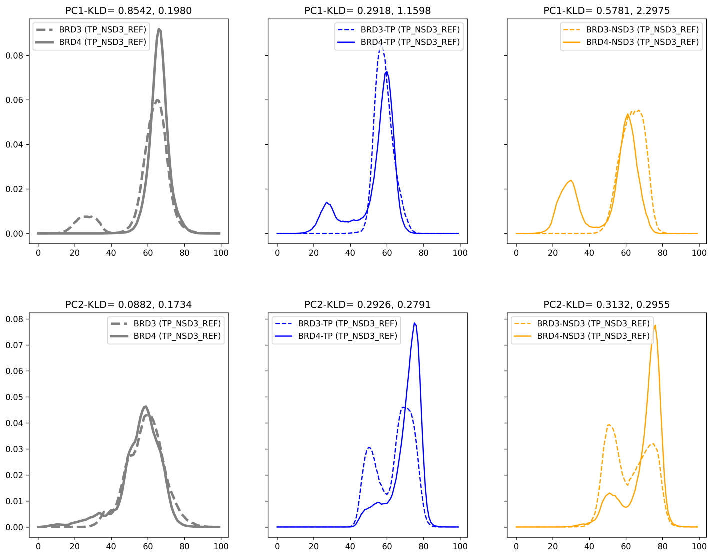


SI Figure 7: Distribution of PC1 (top row) and PC2 (bottom row) and KL divergence values. Comparison between paralogs.

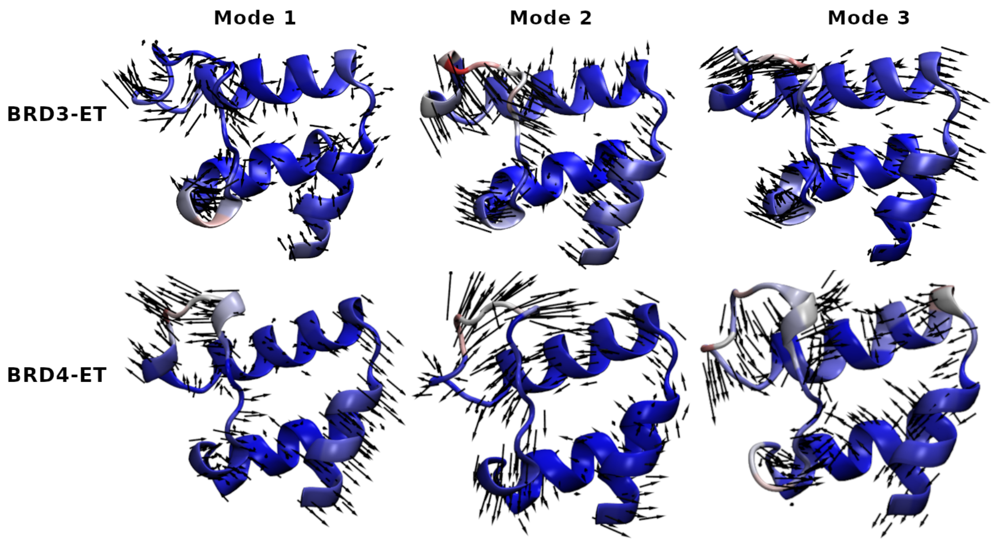


SI Figure 8: Representation of the first three normal modes calculated for BRD3-ET (top) and BRD4-ET (bottom) in their unbound form .

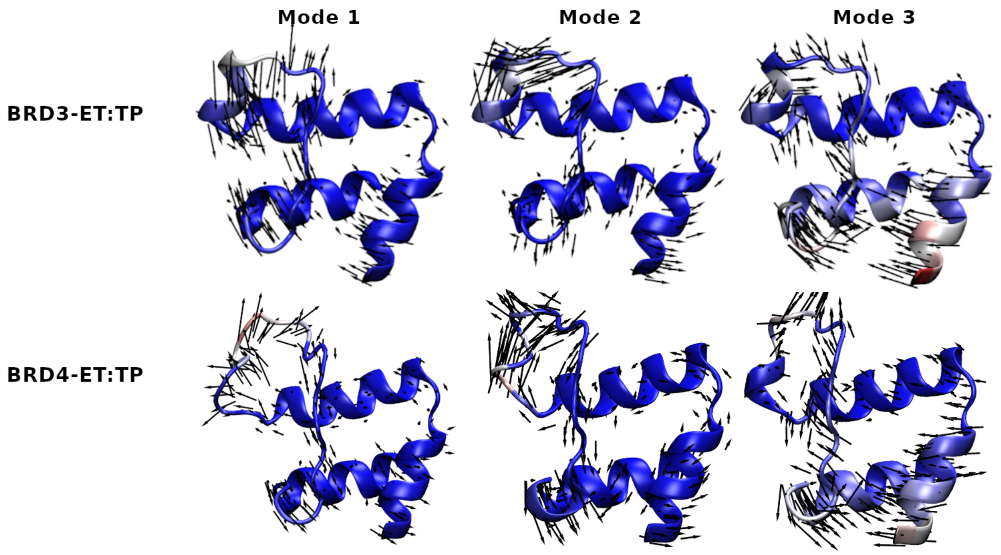


SI Figure 9: Representation of the first three normal modes calculated for BRD3-ET (top) and BRD4-ET (bottom) in their TP-bound form .

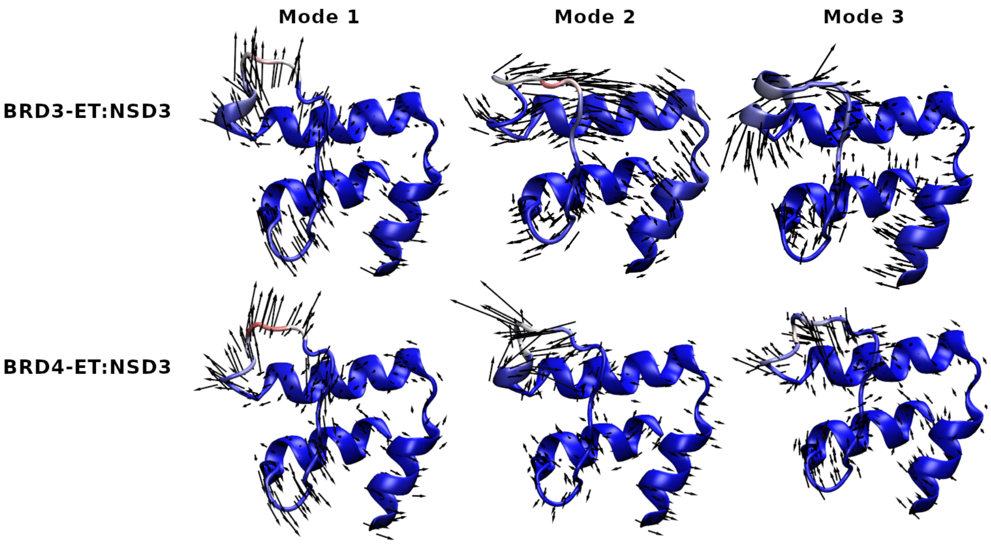


SI Figure 10: Representation of the first three normal modes calculated for BRD3-ET (top) and BRD4-ET (bottom) in their NSD3-bound form .

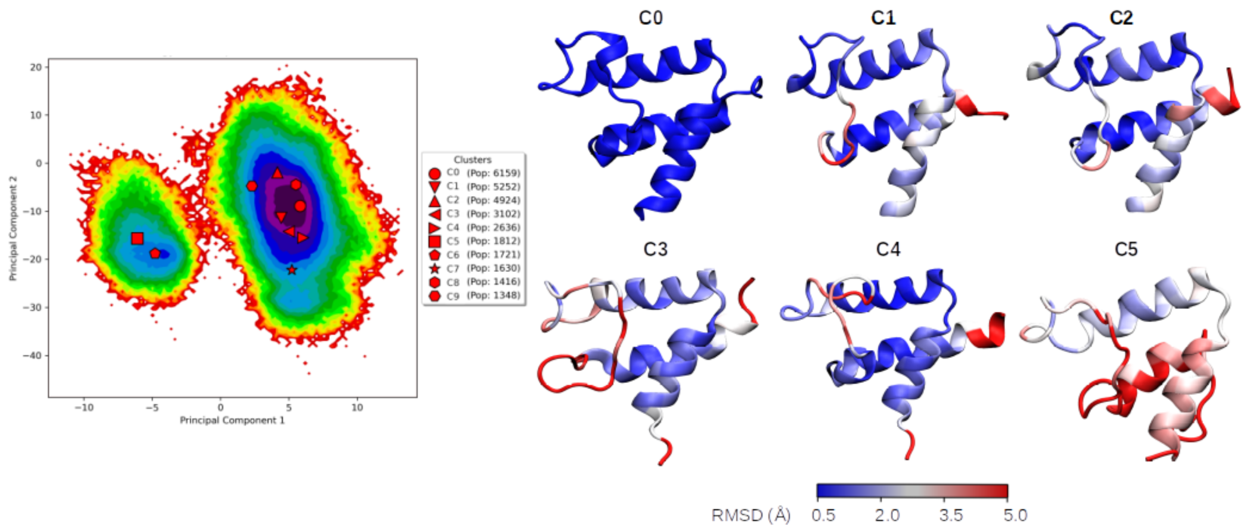


SI Figure 11: Representative structures extracted for BRD3-ET. Structures are colored from blue to red considering the RMSD per residue calculated against the centroid of the top cluster.

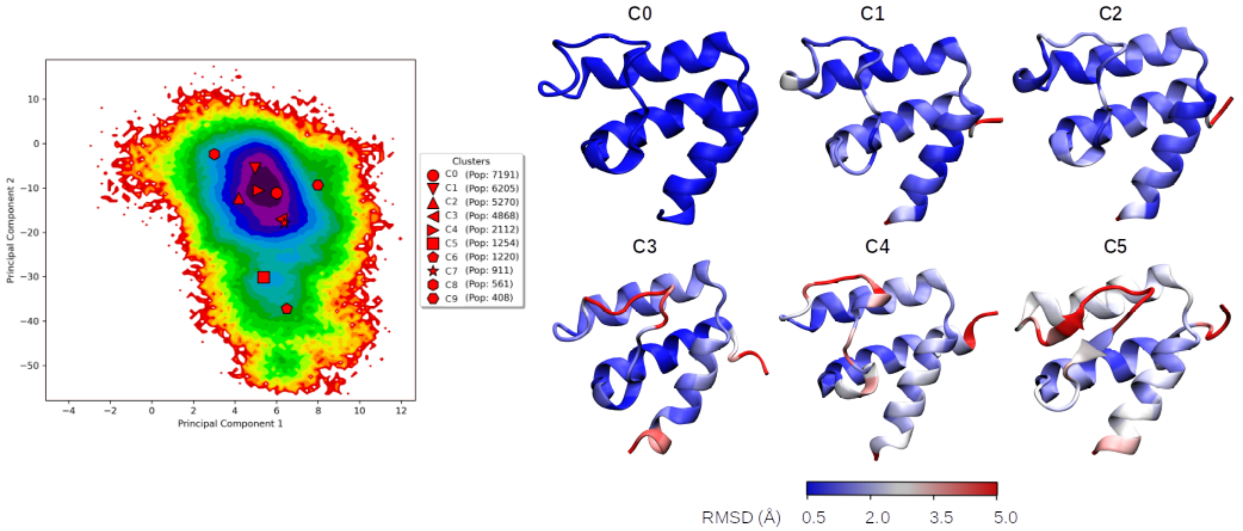


SI Figure 12: Representative structures extracted for BRD4-ET. Structures are colored from blue to red considering the RMSD per residue calculated against the centroid of the top cluster.

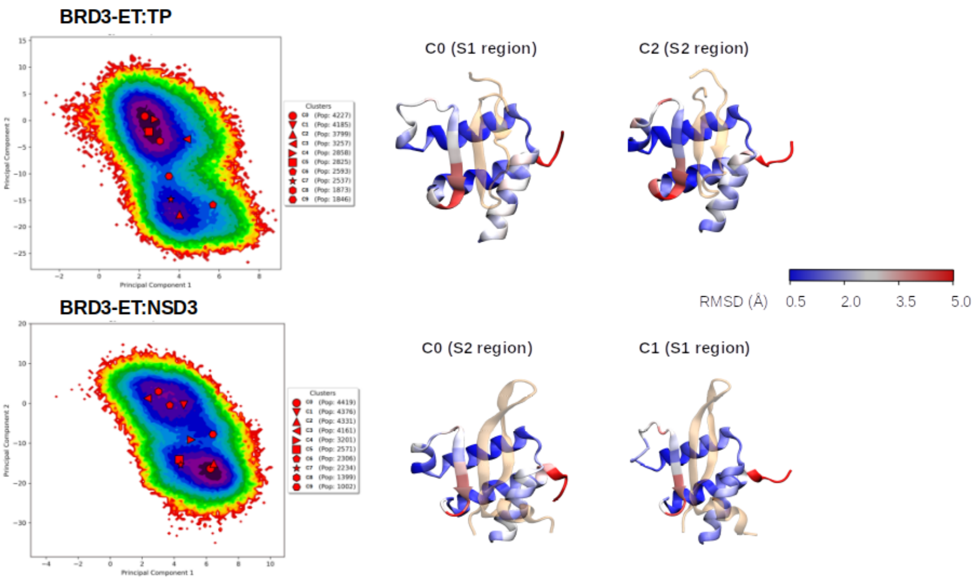


SI Figure 13: Representative structures extracted for BRD3-ET complexes. Structures are colored from blue to red considering the RMSD per residue calculated against the centroid of the top cluster of BRD3-ET unbound. The peptide is shown in color orange with transparency.

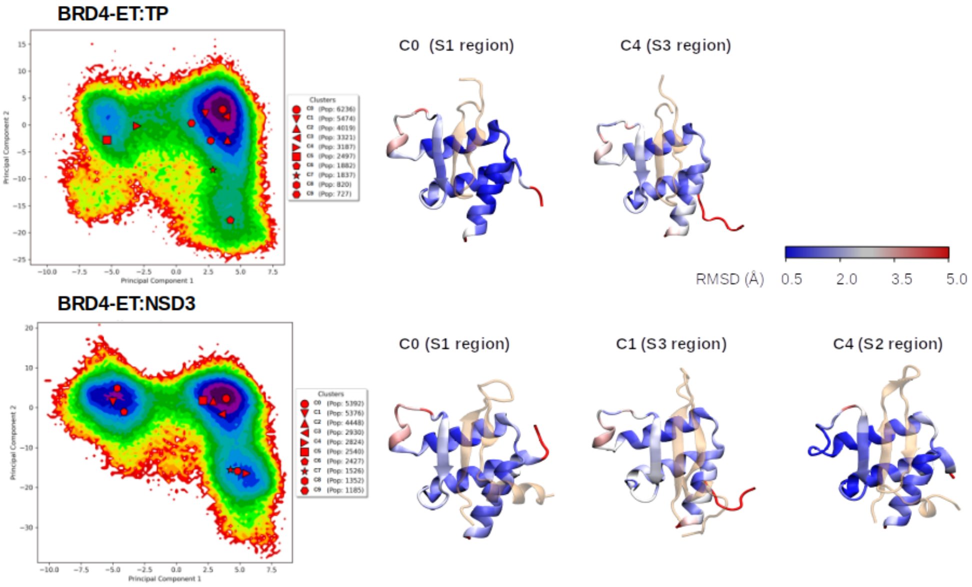


SI Figure 14: Representative structures extracted for BRD4-ET complexes. Structures are colored from blue to red considering the RMSD per residue calculated against the centroid of the top cluster of BRD4-ET unbound. The peptide is shown in color orange with transparency.

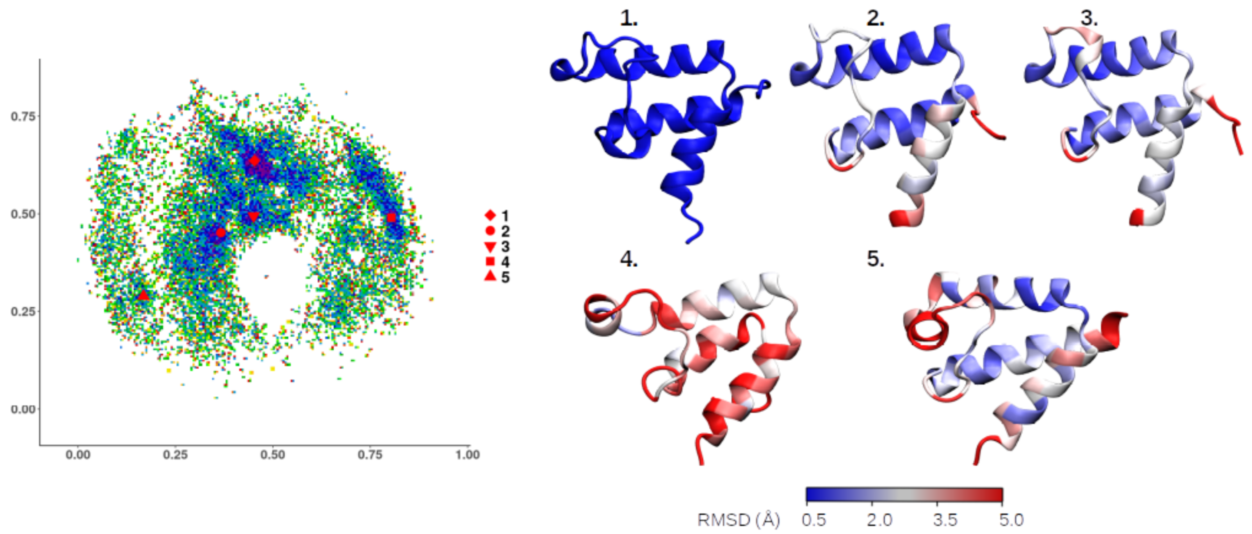


SI Figure 15: Structures extracted from the ELViM map of BRD3-ET. Structures are colored from blue to red considering the RMSD per residue calculated against structure label as 1.

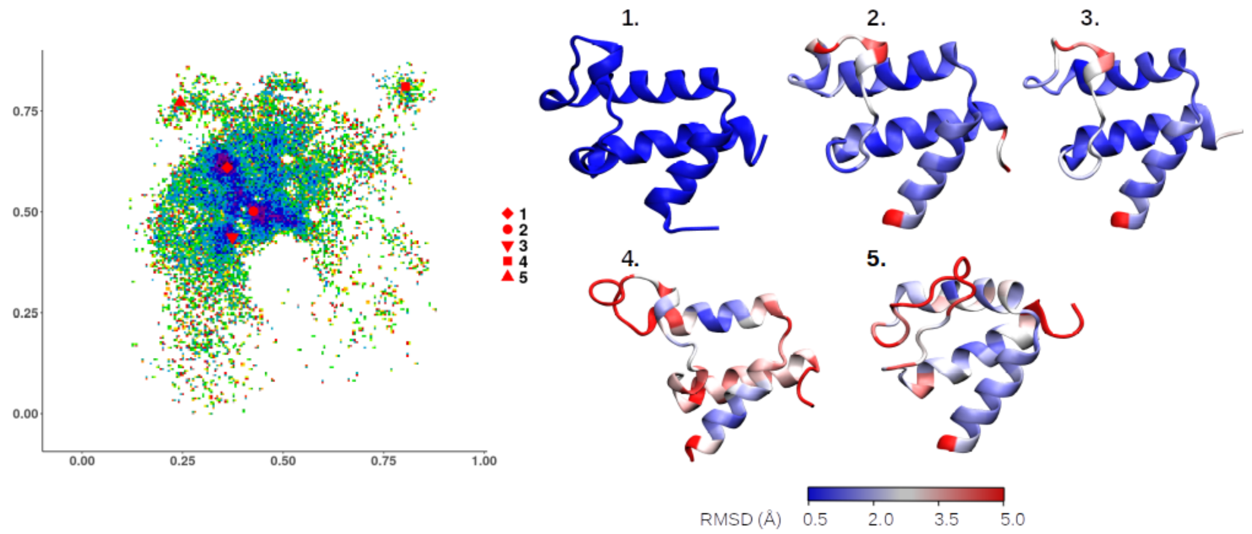


SI Figure 16: Structures extracted from the ELViM map of BRD4-ET. Structures are colored from blue to red considering the RMSD per residue calculated against structure label as 1.

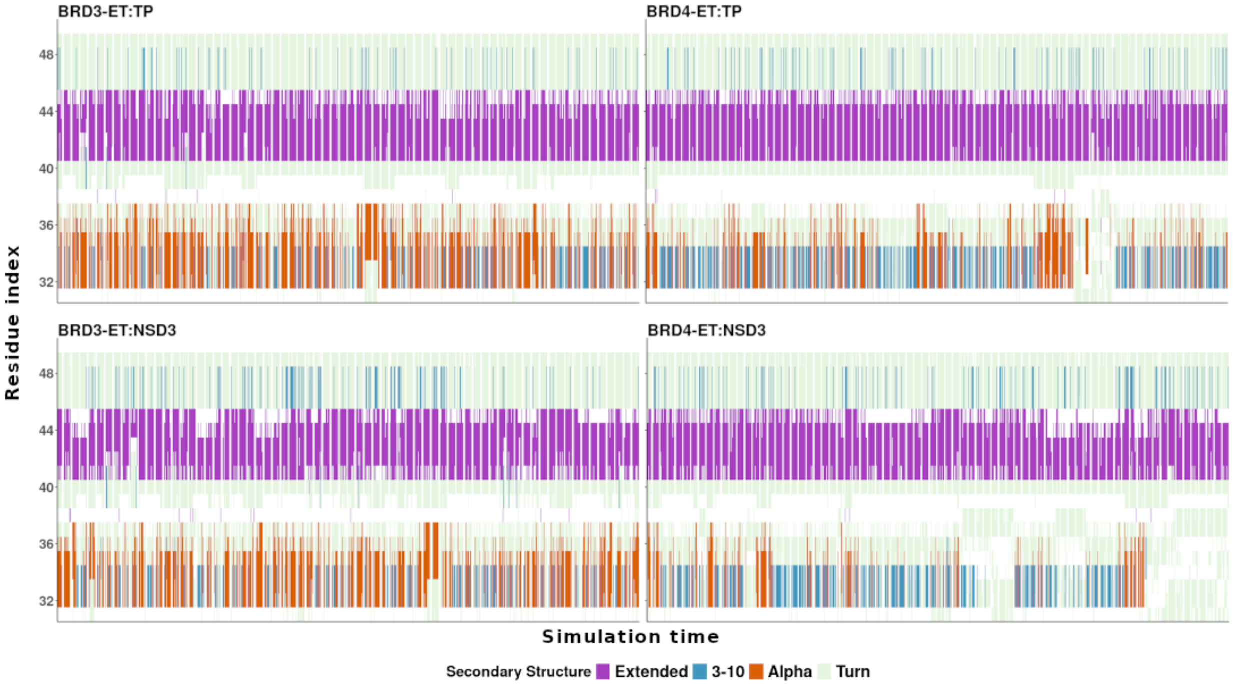


SI Figure 17: Secondary structure content of the loop region as function of the simulation time form the MD, for the bounded forms of BRD3 and BRD4-ET.

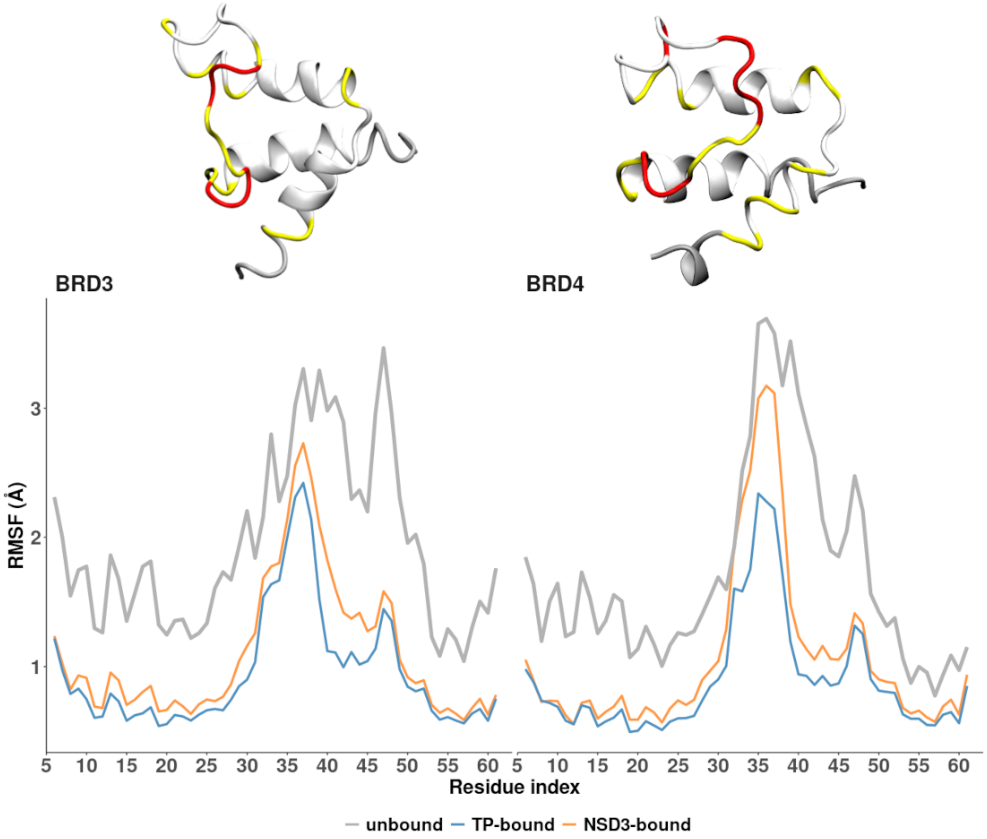


SI Figure 18: RMSF profile per residue. Calculated considering the backbones of residues 6 to 61. The structures highlights the regions that are most stabilized in RMSF terms upon peptide binding with respect to unbound forms. The structures were colored considering the difference between the average RMSF of the bound forms and the unbound form. Yellow: for the residues with $-1.5 \leq \Delta RMSF < -1.0$. Red: residues with $\Delta RMSF < -1.5$.

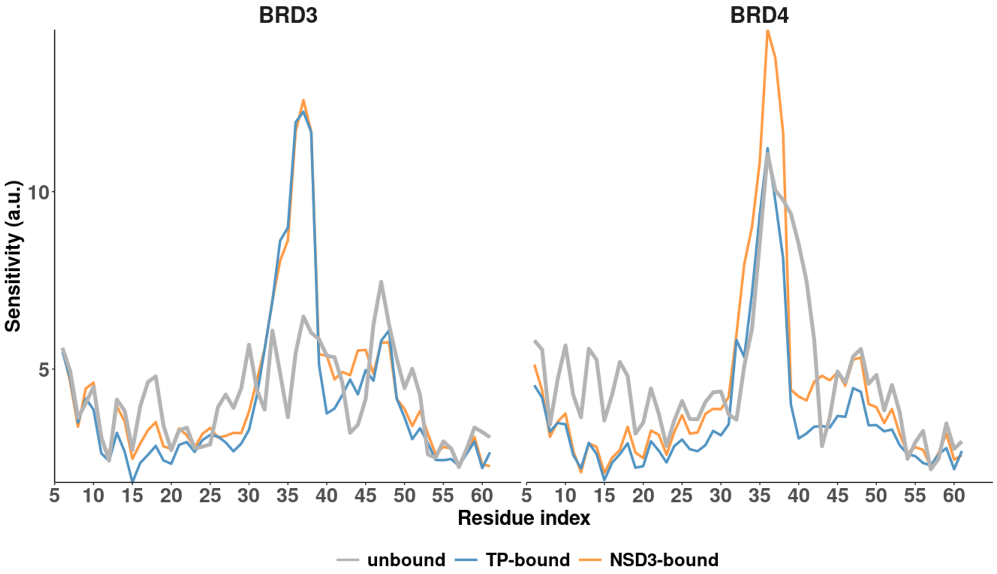


SI Figure 19: Sensitivity profile. Sensitivity is presented in arbitrary units against the residue index. The profile was obtained as the square root of the summation of the column elements of the PRS matrix. The matrix was calculated from the covariance matrix of each system, obtained from the simulations, considering the $C_{alpha}$ of residues 6 to 61 and applying 500 random forces to each site.

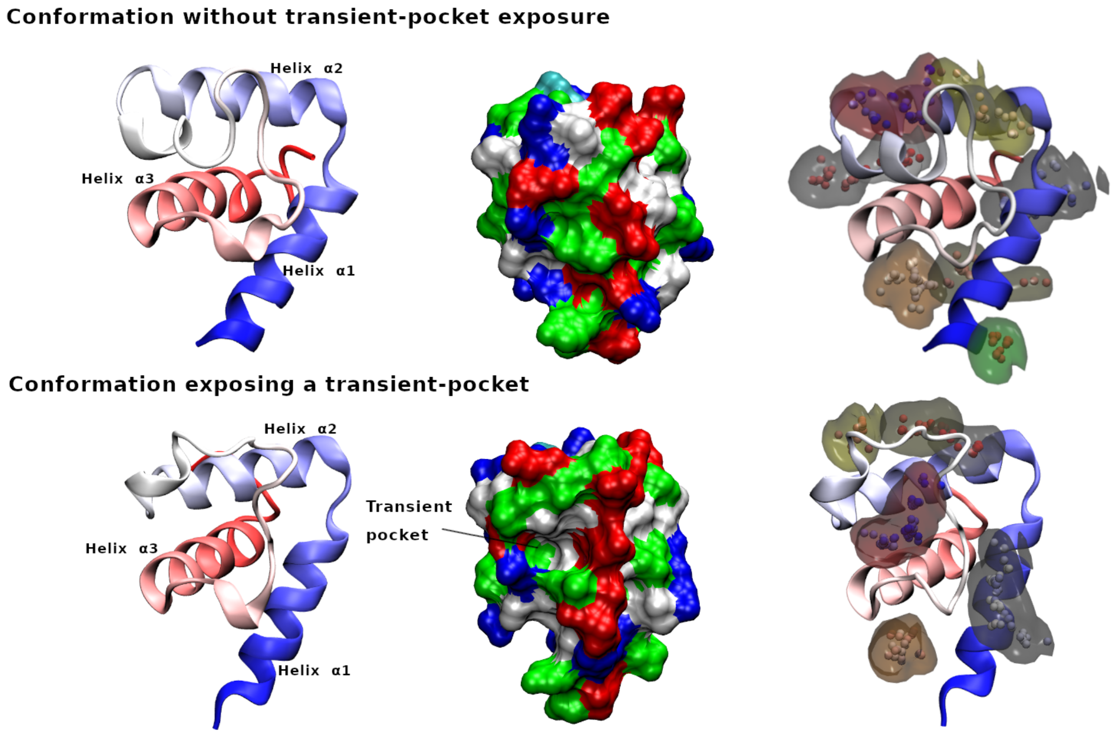


SI Figure 20: Structural comparison showing a transient pocket reveled during the MD simulations. On top we show a conformation without the exposure of the transient pocket (frame 1) and on the bottom, a conformation where the pocket is expose (frame 144772). Representation at the left are colored by residue index (from blue to red) and surface representations in the middle, are colored by residue type. Structures at the right represents the fpocket predictions.

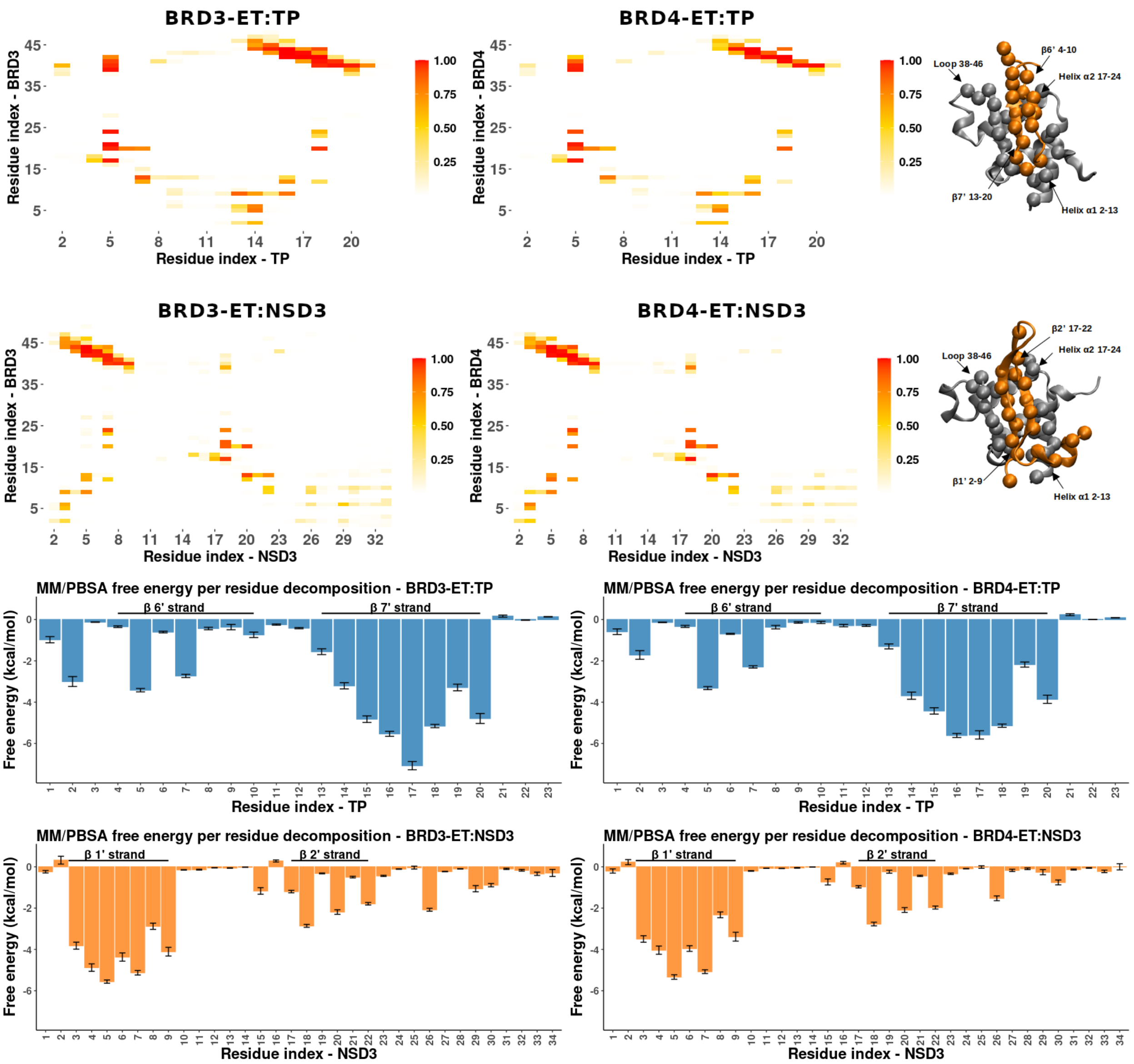


SI Figure 21: Analysis of the protein-peptide interactions. Top: Contact Map between protein receptor and peptide and representation of the structure, highlighting the regions with most frequent contacts. Distance cutoff of 5 Å. The map is colored by contact frequency along the MD trajectories. Bottom: Per-residue MM/PBSA free energy per decomposition. The calculation was performed over 100 frames of the last $\mu s$ of BRD3 and BRD4-ET TP bound and NSD3 bound simulation respectively. Each bar represents the average total free energy variation for each peptide residue and the error bars correspond to the standard error of the mean.

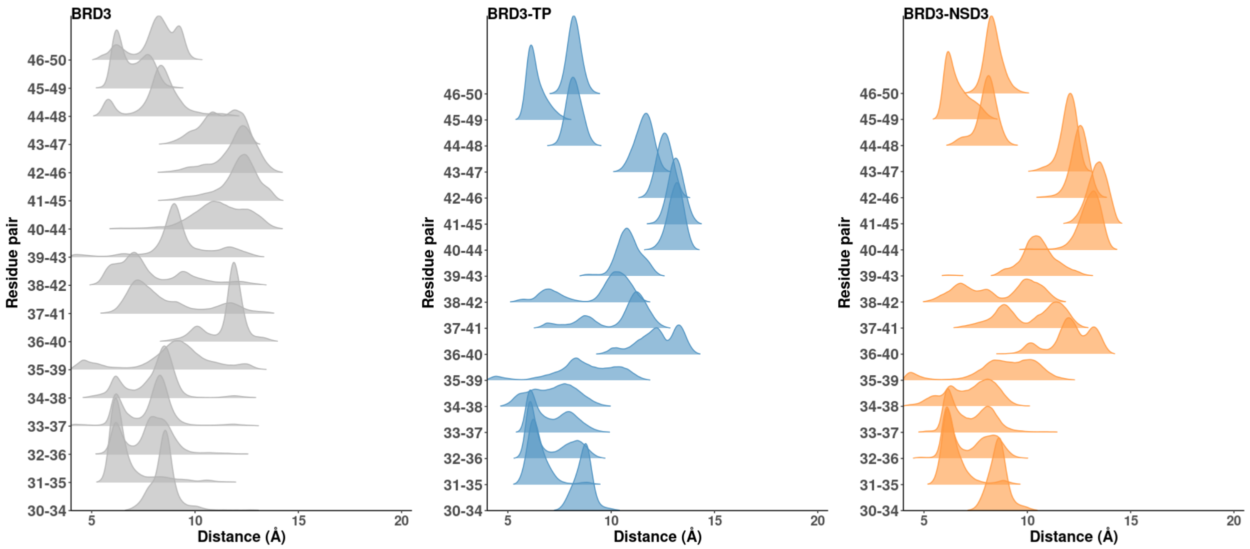


SI Figure 22: Distances between the $C_{alpha}$ of loop residues $i \rightarrow i+4$ for BRD3-ET unbound and bounded forms.

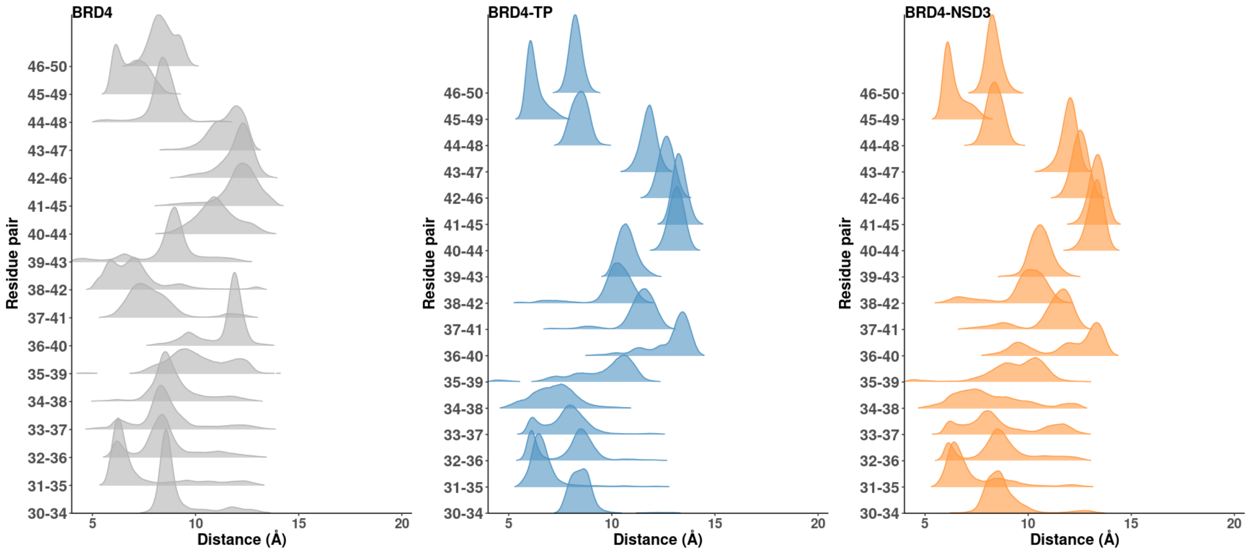


SI Figure 23: Distances between the $C_{alpha}$ of loop residues $i \rightarrow i+4$ for BRD4-ET unbound and bounded forms.

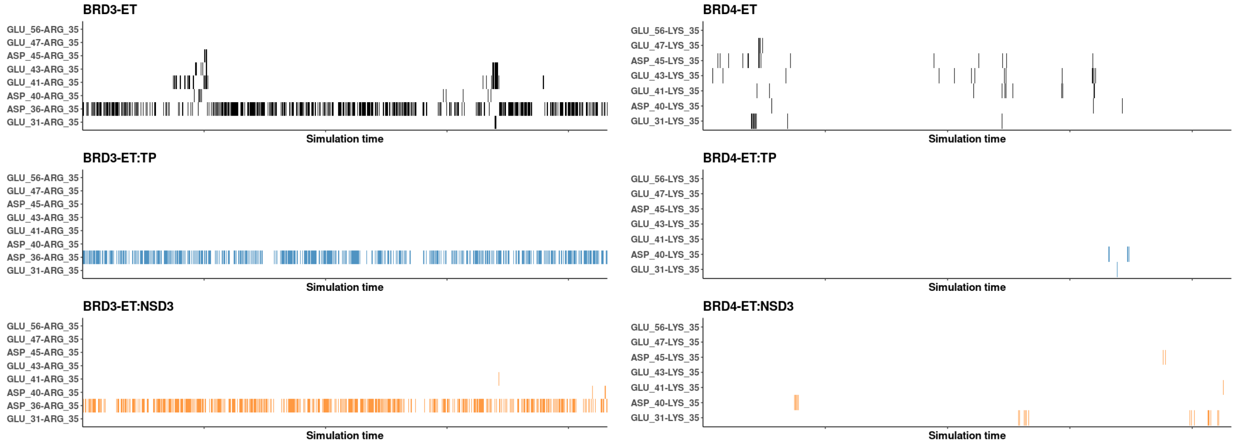


SI Figure 24: Time evolution of salt bridges. The bridges were computed for charged residues withing the receptor considering a distance cutoff of 3.5 Å. The color indicates the presence of the interaction, while its absence is shown in white.

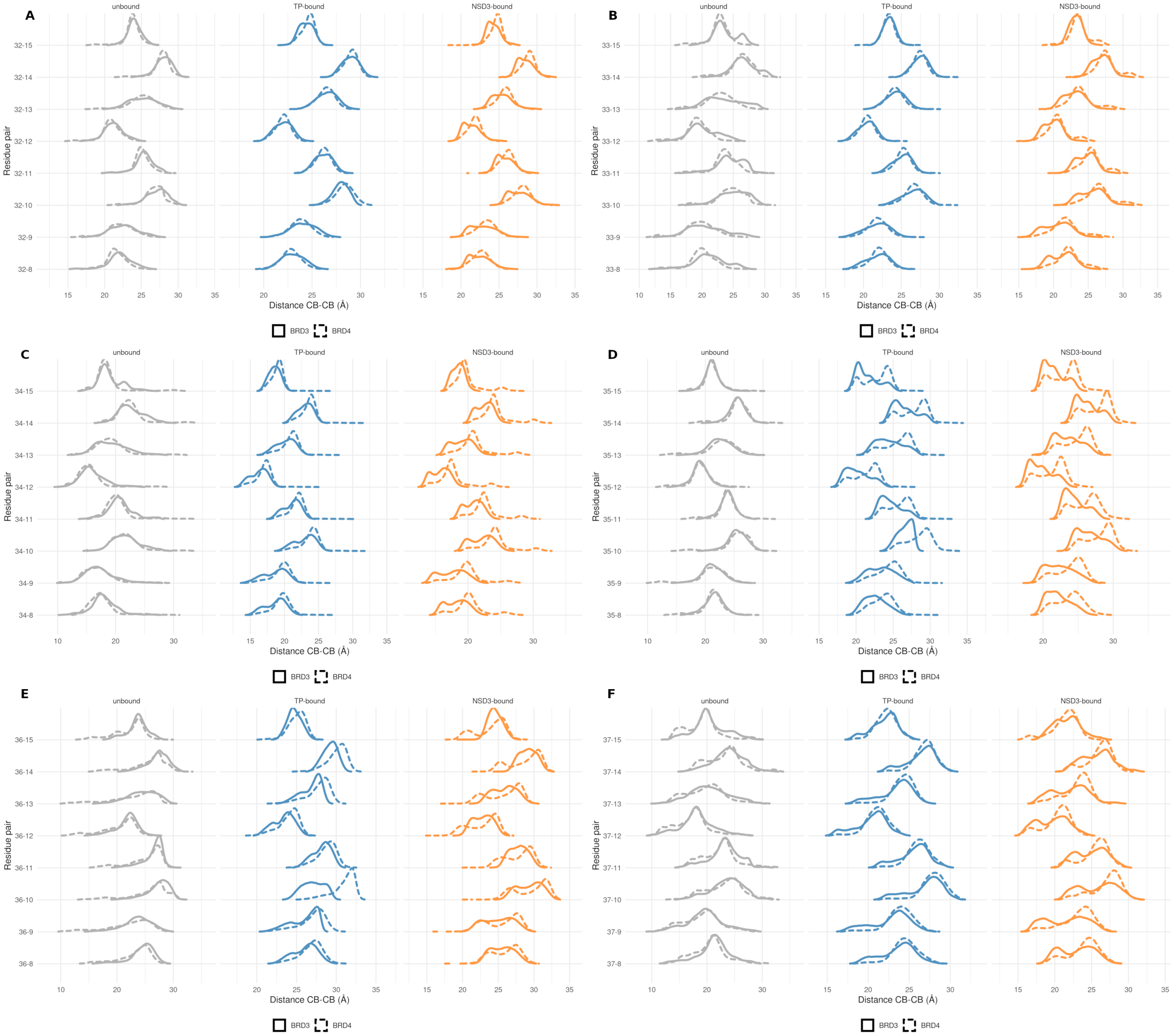


SI Figure 25: Distances between $C_{beta}$ of selected loop residues that conform the $\eta$1-helix (from A to F: residues 32, 33, 34, 35, 36, 37 respectively) and $\alpha$1-helix residues.

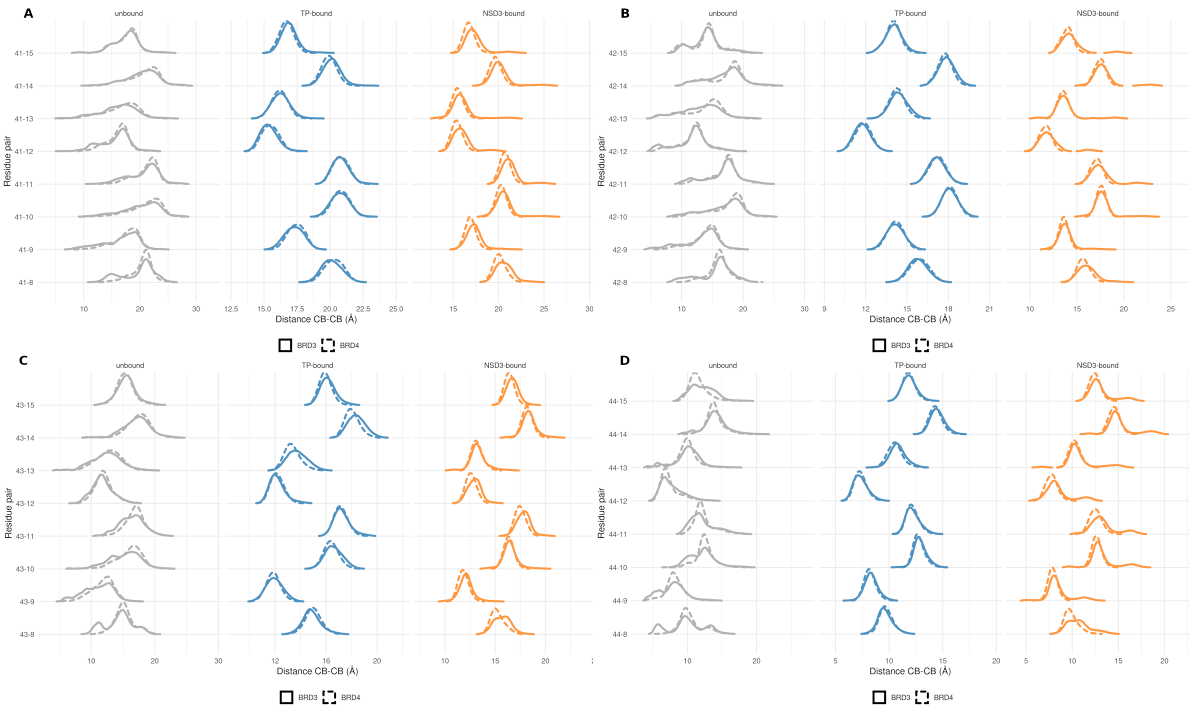


SI Figure 26: Distances between $C_{beta}$ of selected loop residues that conform the $\beta$ strand upon peptide binding (from A to D: residues 41, 42, 43, 44 respectively) and $\alpha$1-helix residues.

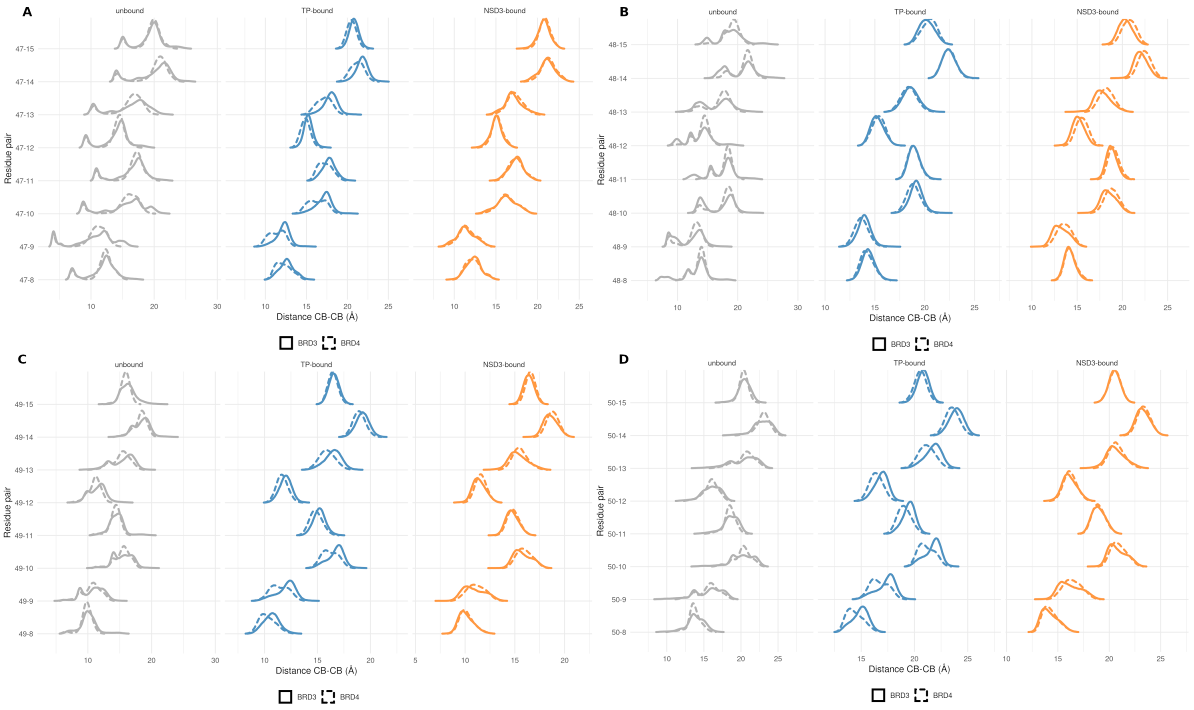


SI Figure 27: Distances between $C_{beta}$ of selected loop residues that conform to the $\eta$2-helix (from A to D: residues 47, 48, 49, 50, respectively) and $\alpha$1-helix residues.
